# Genome scale modeling of the protein secretory pathway reveals novel targets for improved recombinant protein production in yeast

**DOI:** 10.1101/2021.10.16.464630

**Authors:** Feiran Li, Yu Chen, Qi Qi, Yanyan Wang, Le Yuan, Mingtao Huang, Ibrahim E. Elsemman, Amir Feizi, Eduard J Kerkhoven, Jens Nielsen

**Author notes:** These authors contributed equally: Yu Chen, Qi Qi and Yanyan Wang.

## Abstract

Eukaryal cells are used for the production of many recombinant pharmaceutical proteins, including several of the current top-selling products. The protein secretory pathway in eukaryal cells is complex and involves many different processes such as post-translational modifications, translocation, and folding. Furthermore, recombinant protein production competes with native secretory proteins for the limited energy and proteome resources allocated to the protein secretory pathway. Due to the complexity of this pathway, improvement through metabolic engineering has traditionally been relatively ad-hoc; and considering the industrial importance of this pathway, there is a need for more systematic approaches for novel design principles. Here, we present the first proteome-constrained genome-scale protein secretory model of a eukaryal cell, namely for the yeast *Saccharomyces cerevisiae* (pcSecYeast). The model contains all key processes of this pathway, i.e., protein translation, modification, and degradation coupled with metabolism. The model can capture delicate phenotypic changes such as the switch in the use of specific glucose transporters in response to changing extracellular glucose concentration. Furthermore, the model can also simulate the effects of protein misfolding on cellular growth, suggesting that retro-translocation of misfolded proteins contributes to protein retention in the Endoplasmic reticulum (ER). We used pcSecYeast to simulate various recombinant proteins production and identified overexpression targets for different recombinant proteins overproduction. We experimentally validated many of the predicted targets for α-amylase production in this study, and the results show that the secretory pathways have more limited capacity than metabolism in terms of protein secretion.

## Introduction

The protein secretory pathway is an important pathway for eukaryal cells. About 30% of native proteins are processed by the secretory pathway in eukaryotes^1^. The secretory pathway spans several different organelles carrying out peptide translocation, folding, ER-associated protein degradation (ERAD), sorting processes as well as different post-translational modifications (PTMs), ensuring proper protein functionality^2^. There are around 200 proteins engaged in the protein secretory pathway in *Saccharomyces cerevisiae*, hence responsible for these functions. The unique modification profile of each secretory protein dictates specific combinations of multiple processes required for their production and secretion, which makes the secretory pathway a complicated production line and therefore complex to describe. Unraveling the processing and energetic costs for proteins passing through the secretory pathway and how the cell distributes energy and enzymes to process these proteins is therefore desirable, as this would facilitate a better understanding of protein secretion.

*S. cerevisiae* is used as the expression system for around 15% of all protein-based biopharmaceuticals for human use on the market^3^. It has also been used as an important model organism for studying this important pathway, and many discoveries made in yeast translate directly to other eukaryotes such as Chinese Hamster Ovary (CHO) cells that are also widely used for the production of protein-based biopharmaceuticals^4,5^. Since the early days of recombinant protein production in the 1980s, there have been many attempts to improve the protein expression and secretion levels by removing bottlenecks in the protein modification and secretion pathways^6^. However, most of these attempts were usually evaluated for one recombinant protein, and often do not work for improved expression of another protein. Furthermore, the protein yield has typically been much lower than the theoretically estimated range^7,8^. There is therefore much interest in developing a rational design tool for optimization of the secretory pathway for any recombinant protein, in line with what has been developed for metabolism in many cell factories^9,10^.

There are several published frameworks or models for describing protein secretion in yeast and other eukaryotes, but they are either not able to perform simulations or contain only a partial description of the protein secretion pathway^2,11–14^. Besides, even for a recently published secretory model for mammalian cells, the model is solely a basic extension of a genome-scale metabolic model (GEM), which is not able to simulate how native secretory proteins compete with recombinant proteins targeted to pass through this pathway^13^. We, therefore, reconstructed a detailed proteome-constrained genome-scale protein secretory model for *S. cerevisiae* (pcSecYeast), which contains the description of the complete protein secretion pathway and can perform multiple kinds of simulations including the competition of recombinant proteins with native secretory proteins. The model also enables calculation of the energetic cost for native secretory proteins and hereby investigates how misfolded proteins cause growth reduction. We used the model to evaluate the secretion of various recombinant proteins and identify engineering targets for improving their production. The model represents a significant advancement in terms of enabling the more rational design of yeast cells to be used for recombinant protein production, but it also provides a scaffold for building similar models for other eukaryal cells, e.g., CHO cells.

## Results

### Construction of pcSecYeast

We first updated the latest yeast GEM Yeast8^15^ by adding several reactions to enable the synthesis of precursors required in the secretory pathway such as glycosylphosphatidylinositol (GPI) anchor and glycans (Supplementary Table 1). Similar to the metabolic-expression (ME) model for *Escherichia coli*^16^ and *S. cerevisiae*^17^, protein expression, translation, folding, and degradation were then added for all proteins in the model. Besides that, for proteins processed in the secretory pathway, we added reactions detailly describing protein processing including translocation, PTM, folding, misfolding and degradation (Fig. 1a). Hereby the model can describe detailed processes from nascent peptides in the cytosol to the final mature form of proteins in their destination compartment for all proteins in the model. Therefore, pcSecYeast adds a much more detailed description of protein translocation and processing compared with those ME models. To our knowledge, pcSecYeast represents the first model to describe close links between metabolism, protein translation, post-translational protein processing, protein degradation, and protein secretion in yeast and can be easily adapted to any cell. The components that participate in the protein secretory pathway are involved in 12 subsystems (Fig. 1b). Overall, pcSecYeast accounts for 1,639 protein-coding genes and approximately 70% of the total proteome mass according to PaxDb^18^ (Supplementary Table 2). Details of the reconstruction process can be found in the Supplementary Methods.

**Fig. 1:**
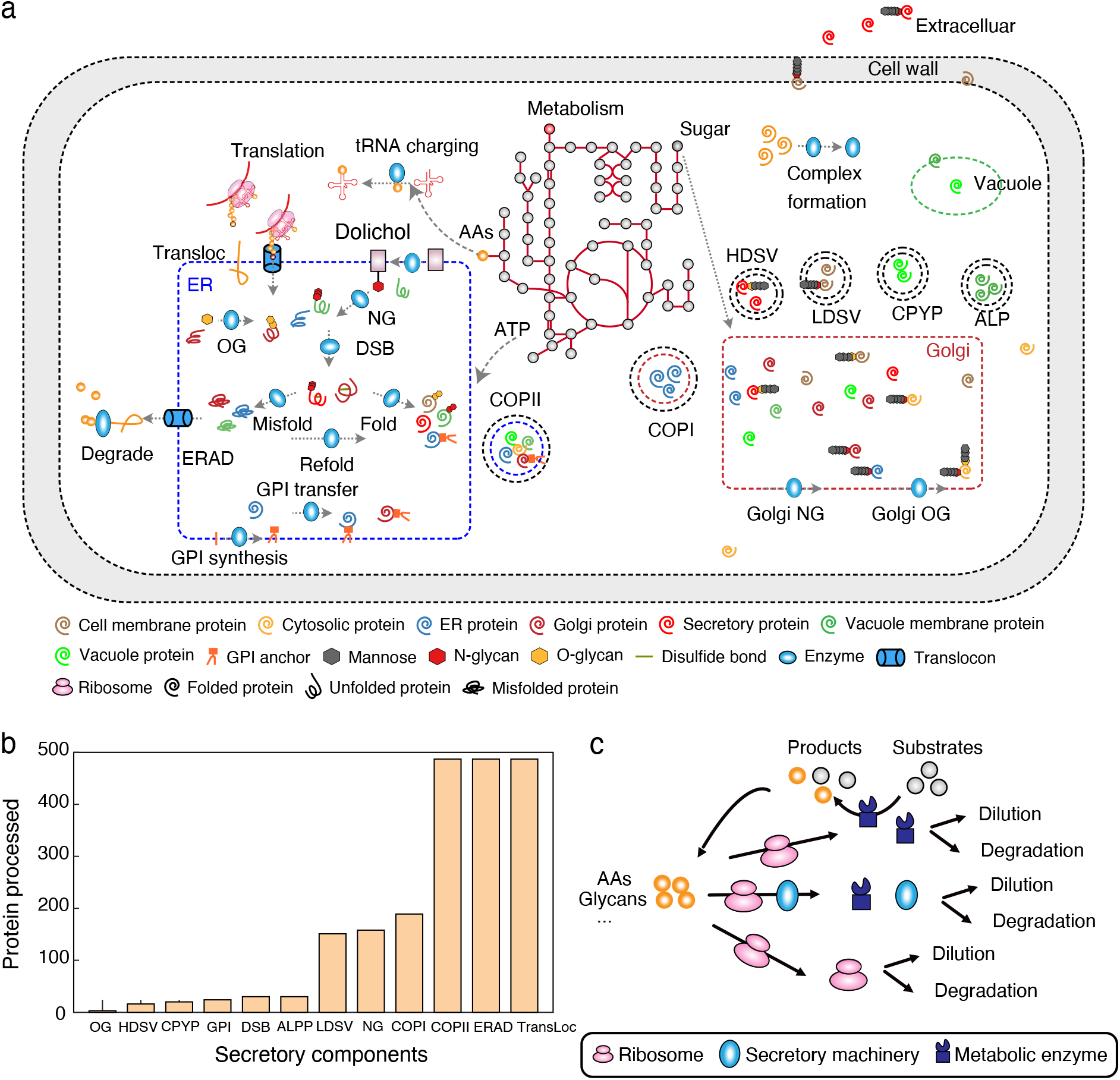
Overview of components in pcSecYeast. a) Simplified schematic processes involved in the protein secretion pathway. The process includes protein translation, translocation, glycosylate, GPI transfer, ERAD and sorting process. The detailed description of all components and reactions can be found in Supplementary Methods. Transloc: translocation, NG: N-glycosylation, OG: O-glycosylation, DSB: disulfide bond formation, GPI: glycosylphosphatidylinositol, ER: endoplasmic reticulum, ERAD: ER-associated degradation, LDSV: low-density secretory vesicles, HDSV: high-density secretory vesicles, ALPP: alkaline phosphatase pathway, CPYP: carboxypeptidase Y pathway. b) Subsystems in the secretory pathway and the number of proteins that are processed in each subsystem. c) Coupling process in the model. Metabolic part produces energy and precursors such as amino acids, glycans for enzyme and ribosome synthesis. Enzymes constrain these metabolic reactions. Ribosomes constrain protein translation. The secretory machinery constrains protein processing in this pathway. All proteins, including ribosomes are diluted due to growth and degraded due to misfolding.

As an extension of Yeast8, pcSecYeast includes default constraints such as mass conservation and flux bounds on metabolic reactions. Besides them, we introduced coupling constraints to relate protein synthesis with metabolism (Supplementary Methods). The metabolic part in the model supplies the substrate and energy for the protein-related part such as ribosome and enzyme synthesis, while the metabolite conversion process in the metabolic part is catalyzed by enzyme complexes synthesized in the protein-related part (Fig. 1c). Protein synthesis is constrained by the synthesis of ribosome and other machineries such as secretory machinery complexes (Fig. 1c). Each flux of enzymatic reaction in the model is constrained at the maximal rate of the associated enzyme, which is a function of turnover rate (*k*_cat_) and the enzyme concentration. Thus, we can simulate the minimum protein levels which sustain the metabolic state, i.e., the proteome-constrained metabolic state. This means that the proteome composition in pcSecYeast is not a fixed amount of average amino acid compositions as in the default GEMs, but a dynamic changing composition of enzymes which reflects the cell state at a certain condition. Thus, the model enables simulation of resource allocations in the cell under different conditions, such as how the cell would balance recombinant protein with native secretory proteins in the recombinant protein production and how the cell would optimize its enzyme profile among various environmental conditions.

### Secretory cost initiates the switch of hexose transporters

Transporters are one important group of proteins that pass through the secretory pathway. Yeast has multiple hexose transporters with diverse kinetics, which are expressed at different levels under different extracellular glucose concentrations^19^. To investigate how the model can simulate the expression and processing of glucose transporters, we utilized the model to simulate yeast growth under different glucose concentrations (Methods). As a result, the model captured the metabolic shift referred as the Crabtree effect, i.e., the production of ethanol at high specific growth rates (Fig. 2a). Furthermore, the model correctly predicted a switch from the predominant use of the high-affinity glucose transporter (Hxt7) to low-affinity glucose transporters (Hxt3 and Hxt1) at high glucose concentrations (Fig. 2b), which is consistent with the experimental observation that *HXT3* and *HXT1* genes are only expressed at high specific growth rates^19^. This is explained by the difference in kinetics of the different sugar transporters, i.e., *k*_cat_ and *K*_M_, and therefore the secretory cost for synthesizing and processing glucose transporters that can support a given glucose uptake flux. Thus, at low specific growth rates where there is a low glucose uptake rate, the cells express a high-affinity transporter with a low *k*_cat_ in a small amount, but to support a high glucose uptake rate it is necessary to express a large amount of glucose transporters, then the low-affinity transporters with high *k*_cat_ values are preferred. This is illustrated by eq. 1, which specifies the secretory cost for a glucose transporter to sustain a given glucose uptake rate can be calculated as the ‘unit secretory cost’ multiplied by the abundance. The protein abundance of the transporter [E_*i*_] is determined by the glucose uptake rate *V*_*glc*_ *K*_M_ and extracellular glucose concentration [*S*] according to the Michaelis-Menten equation. The ‘unit secretory cost’ is defined as the cost required for translation, modification, and secretion of one mol specific protein, which can be predicted by pcSecYeast (Methods). We predicted the ‘unit secretory costs’ for all native secretory proteins in *S. cerevisiae* (Supplementary Table 3) and found that Hxt1 and Hxt3 have a smaller ‘unit secretory cost’ compared with Hxt7, suggesting that synthesizing one mol Hxt1 and Hxt3 would pose less energy burden on the cell. This is partly because Hxt1 has fewer N-glycosylation modification sites than Hxt7 (Supplementary Table 4). Combining with the glucose uptake rate, extracellular glucose concertation, *k*_cat_, and *K*_M_, we can calculate the secretory cost for each glucose transporter from the eq.1 (Fig. 2c). The result suggests that with an increase in the glucose concentration, utilization of Hxt1 and Hxt3 would gradually gain the advantage over Hxt7 (Fig. 2c). Parameter sensitivity analysis of Hxt1 showed that even if we set the same *k*_cat_ for Hxt1 and Hxt7, Hxt1 would still be favorable for glucose uptake in the model simulation at maximum growth rate (Supplementary Figure 1). This demonstrates that the contribution of the low ‘unit secretory cost’ of Hxt1 is critical. Our model hereby predicts that the switch of different affinity glucose transporter is a resource optimization strategy of the cell to adapt to limited resources.

**Fig. 2:**
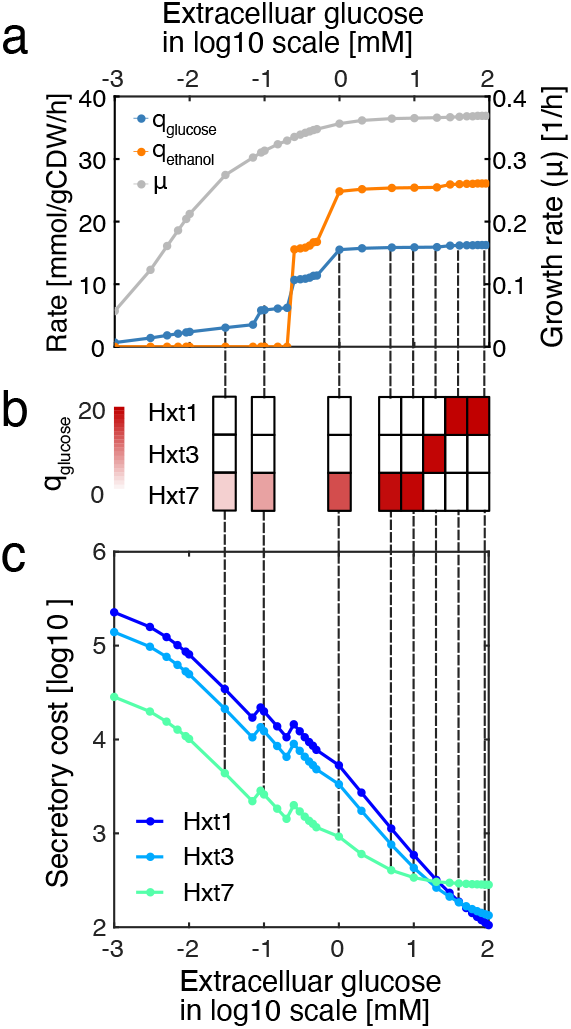
Simulated physiological response of *S. cerevisiae* as a function of the extracellular glucose concentration. a) Simulated glucose uptake rates, ethanol production rates and specific growth rates under different extracellular glucose concentrations. Each point is the simulated result under a certain extracellular glucose condition. b) Specific glucose uptake rate carried by each glucose transporter. Hxt1 and Hxt3 are two low-affinity glucose transporters, while Hxt7 is a high-affinity glucose transporter. c) Calculation of secretory costs of different glucose transporters with the specific glucose uptake rate at input for each extracellular glucose concentration, unit secretory cost, K_M_ and *k*_cat_ that are specific to each transporter based on eq. 1 in the text.

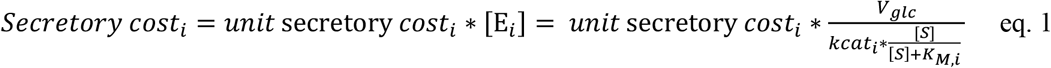

### Yeast suppresses expression of high-cost secretory proteins under secretion pressure

The protein secretory pathway is concurrently processing hundreds of proteins that compete for limited resources such as energy, precursors, and components of the secretory machinery. It has been reported that recombinant mammalian cells repress the expression of native energetically expensive secretory proteins to save limited resources for growth and recombinant protein production^13^. With our proteome allocation model of the secretory pathway, we can perform not only the same calculation of the secretory costs of all 497 native secretory and cell membrane proteins as done for mammalian cells^13^ (denoted as ‘direct cost’ in the Supplementary Figure 2a) but also a more accurate analysis of the costs including the associated costs for corresponding shares of catalyzing enzymes and secretory machineries required for processing the protein besides the cost for itself (‘unit secretory cost’ in Supplementary Figure 2a). By correlating ‘unit secretory cost’ with ‘direct cost’, we found that the ‘unit secretory cost’ calculated in pcSecYeast is overall 3.8-fold higher compared with the ‘direct cost’ (Supplementary Figure 2a). Outliers in the correlation of these two kinds of cost calculation are mainly caused by the unusual protein features such as the 52 N-glycosylation sites annotated for the protein RAX2 or long amino acid sequences for large proteins TOR1 and TOR2 (Supplementary Figure 2a). To evaluate if there is reduced expression of the proteins that are costly to process by the secretory pathway as observed in mammalian cells, we correlated the calculated ‘unit secretory costs’ with the mRNA levels of 497 native secretory proteins for three strains with different levels of recombinant α-amylase production that were characterized in a recent study^20^. We observed a significant negative correlation (*P* value < 1e-8) between unit secretory costs and mRNA levels of native secretory proteins in all three strains (Supplementary Figure 2b-c), suggesting that the cells suppress the expression of proteins that are expansive to secrete when the secretory pathway is under pressure to process a recombinant protein. Moreover, we found that the negative correlations are stronger in the strains with higher α-amylase production levels (MH34 and B184) compared with that in the strain with a lower α-amylase production level (AAC) (Supplementary Figure 2c, *P* value = 0.004). Therefore, the suppression level for costly native secretory proteins depends on the recombinant protein production levels, suggesting that the yeast cells respond accordingly to the level of secretion stress.

### Misfolded protein slows maximum growth

Protein synthesis and secretion is an error-prone process. Mutation in the sequence, errors during the synthesis or environmental insults cause the newly synthesized protein to misfold^21^. Misfolded proteins are prioritized to be eliminated rapidly by the ERAD pathway but may retain and accumulate in the ER, and could trigger cell stress (Fig. 3a)^22–24^. Here, we used our model to simulate the ER tolerance to misfolded proteins. We expanded pcSecYeast to include the production of vacuolar carboxypeptidase Y (YMR297W, CPY), since CPY and its derived misfolded form CPY* are processed in the secretory pathway, and widely used in the elucidation of the mechanisms of ER quality control and ERAD of misfolded proteins^25^. By modifying misfolding ratio parameter in the model, we can simulate the misfolding levels of CPY. A misfolding ratio of 1 means that all the CPY protein molecules are misfolded and cannot be targeted to the Golgi for further processing as the reported misfolded form CPY* in literature^26^.

**Fig. 3:**
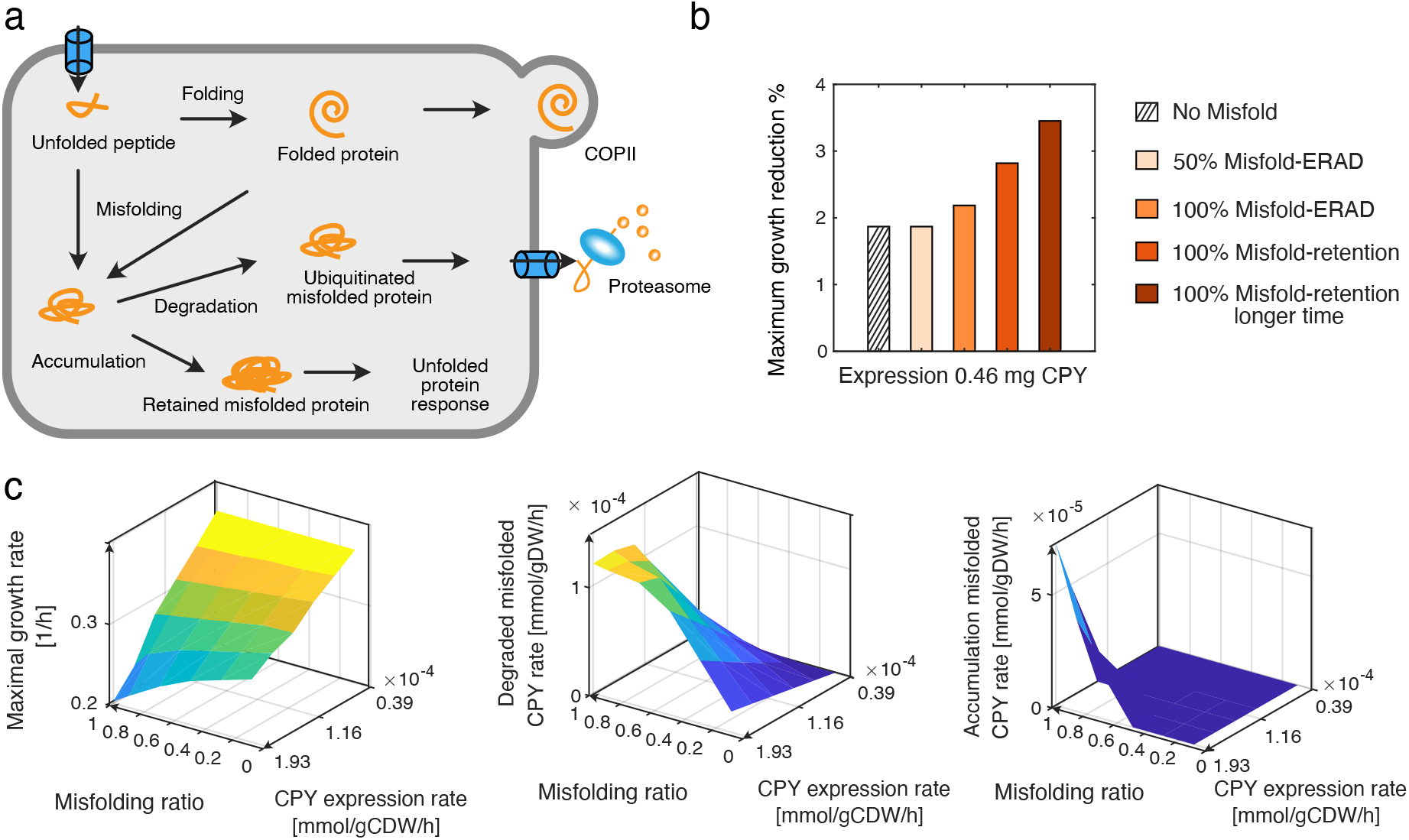
Simulation of CPY overexpression. a) Schematic view of different routes for expressed CPY. b) Reduction of simulated maximum specific growth rate [1/h] due to expression at certain levels of CPY following different routes. c) Simulations for various CPY expression levels and misfolding ratios with the constraint for retro-translocation enzymes.

Here, we used the maximum growth rate reduction to indicate the fitness cost for CPY going through different routes: 1) all correctly folded and targeted to the vacuole without misfolding; 2) misfolded in different ratios and some targeted for ERAD; 3) all of them misfolded, retained in the ER at different times. Our simulations showed that misfolding imposes more fitness cost compared with correct folding; that retention imposes more fitness cost compared with ERAD; and that retention in the ER for a long time would also impose more fitness cost. We predicted that there is about 1.9% maximum specific growth reduction when expressing 0.46 mg native CPY without misfolding (0.46 mg representing 0.1% of the total proteome, Fig. 3b), which is comparable to the measurement for expression of a cytosol wildtype Yellow Fluorescent Protein (YFP) where a 1.4% growth reduction was observed for expressing YFP at the same abundance^27^. Compared with cytosolic YFP, CPY requires extra energy and allocation of resources in the secretory pathway, and this can explain the slightly higher predicted growth reduction for expressing CPY. The growth reduction for expressing CPY proteins all in misfolded form is 2.2%-3.5% (Fig. 3b), which is also comparable with the measurement of expressing the mostly misfolded cytosolic YFP at the same level (up to 3.2% growth reduction)^27^. The growth reduction measured for YFP in the literature is a combination of fitness cost caused by the misfolding itself and unfolded protein response in the cytosol (UPR-cyto) triggered by the accumulation of misfolded proteins. In the model simulations, we only considered the fitness cost for misfolding and degradation of CPY. This also suggests that ER misfolded protein imposes more fitness cost compared with cytosolic misfolded protein when they are expressed at the same level. If the misfolded proteins are degraded by ERAD and the proteasome, then amino acids and modification precursors such as glycans can be recycled. However, if misfolded proteins are retained in the ER, they would compete with unfolded protein for limited ER quality control proteins especially Kar2 and Pdi1^27^, which would further increase the ER burden. We investigated the simulated various protein levels and found that the levels of Kar2 and Pdi1 increase significantly when CPY is retained (Supplementary Figure 3), which suggests that the retained protein would drain Kar2 and Pdi1 and therefore compete with native proteins processed in the secretory pathway. In addition, we evaluated the ER redox stress by comparing the transport of glutathione (GSH) and glutathione disulfide (GSSG) and found that the flux of GSSG export from the ER is significantly higher when misfolded protein is retained in the ER (Supplementary Figure 4), suggesting the higher redox unbalance in the ER at this state.

**Fig. 4:**
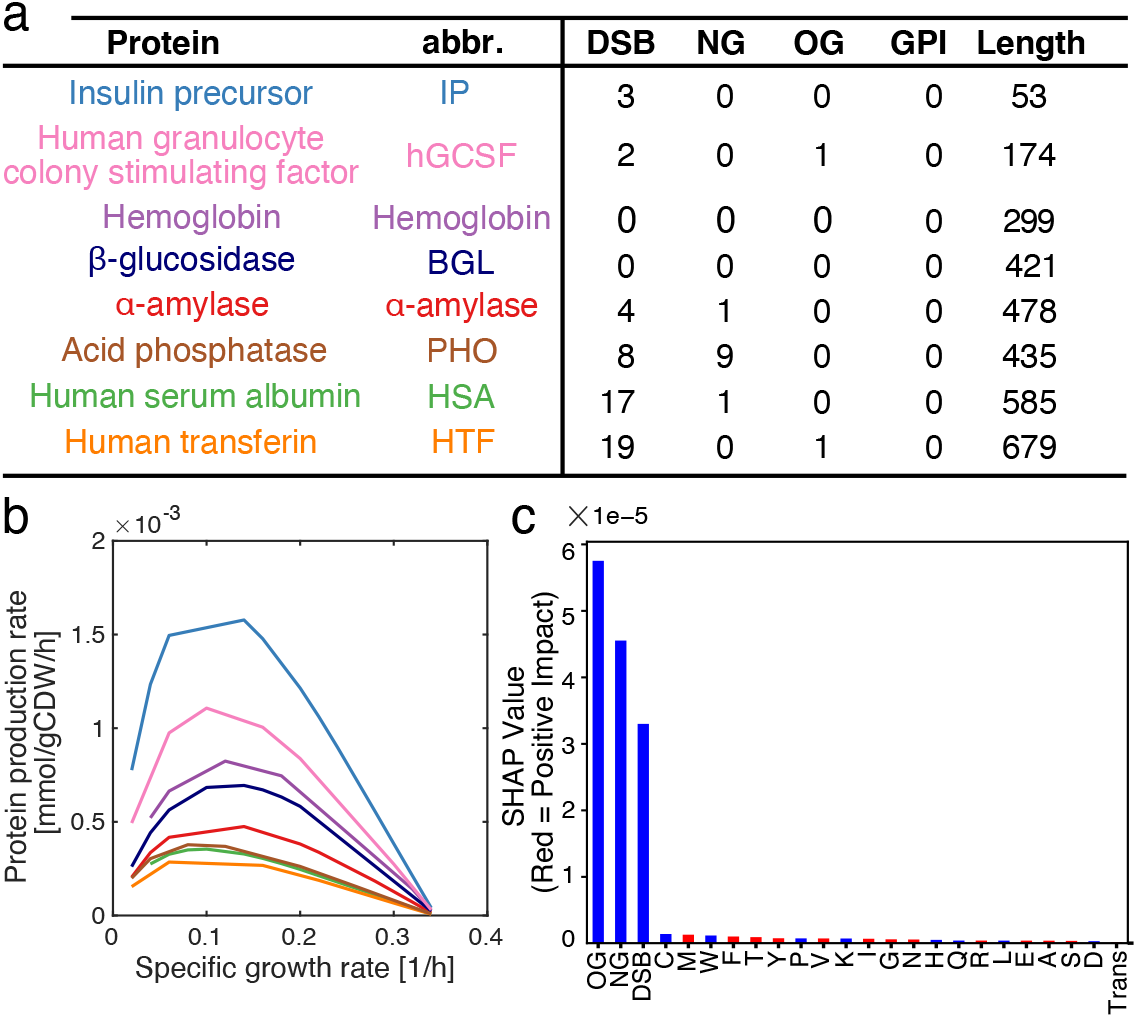
Simulation of recombinant protein production. a) Overview of protein features for eight recombinant proteins produced by *S. cerevisiae*. See Supplementary Table 5 for detailed information. b) Simulation of maximum specific recombinant protein production rate as a function of specific growth rate. c) Feature importance analysis towards recombinant protein production. NG: N-glycosylation site; OG:O-glycosylation site; DSB: disulfide bond number; trans: transmembrane domain; one letter stands for amino acid. Blue color stands for negative impact of having this feature towards recombinant protein production rate, while red color indicates a positive impact.

Furthermore, we performed analysis to identify parameters leading to misfolded protein accumulation in ER (Supplementary Figure 5a-d, Fig. 3c) and found that when retro-translocation enzymes (Sec61, Doa10 and Hrd10) are constrained, the excessive misfolded CPY would be retained and accumulated in ER if CPY is expressed at high levels, causing a steeper decrease in the specific growth rate (Fig. 3c). Other parameters such as ERAD capacity, ER volume, ER membrane space and secretory machinery capacity were not able to show the retention and accumulation phenotype when constrained in the model (Supplementary Figure 5a-d). We found that the retention of the misfolded protein phenotype is alleviated when removing the constraint of retro-translocation enzymes, suggesting the importance of the retro-translocation towards handling of misfolded proteins (Supplementary Figure 5e). Therefore, we can use the pcSecYeast model with the extra constraint of retro-translocation enzymes to mimic state of misfolded protein accumulation in ER (Fig. 3c). The plateau in the CPY degradation rate demonstrates that there is a maximum capacity of the ERAD pathway and therefore also a tolerance limit for misfolded CPY.

**Fig. 5:**
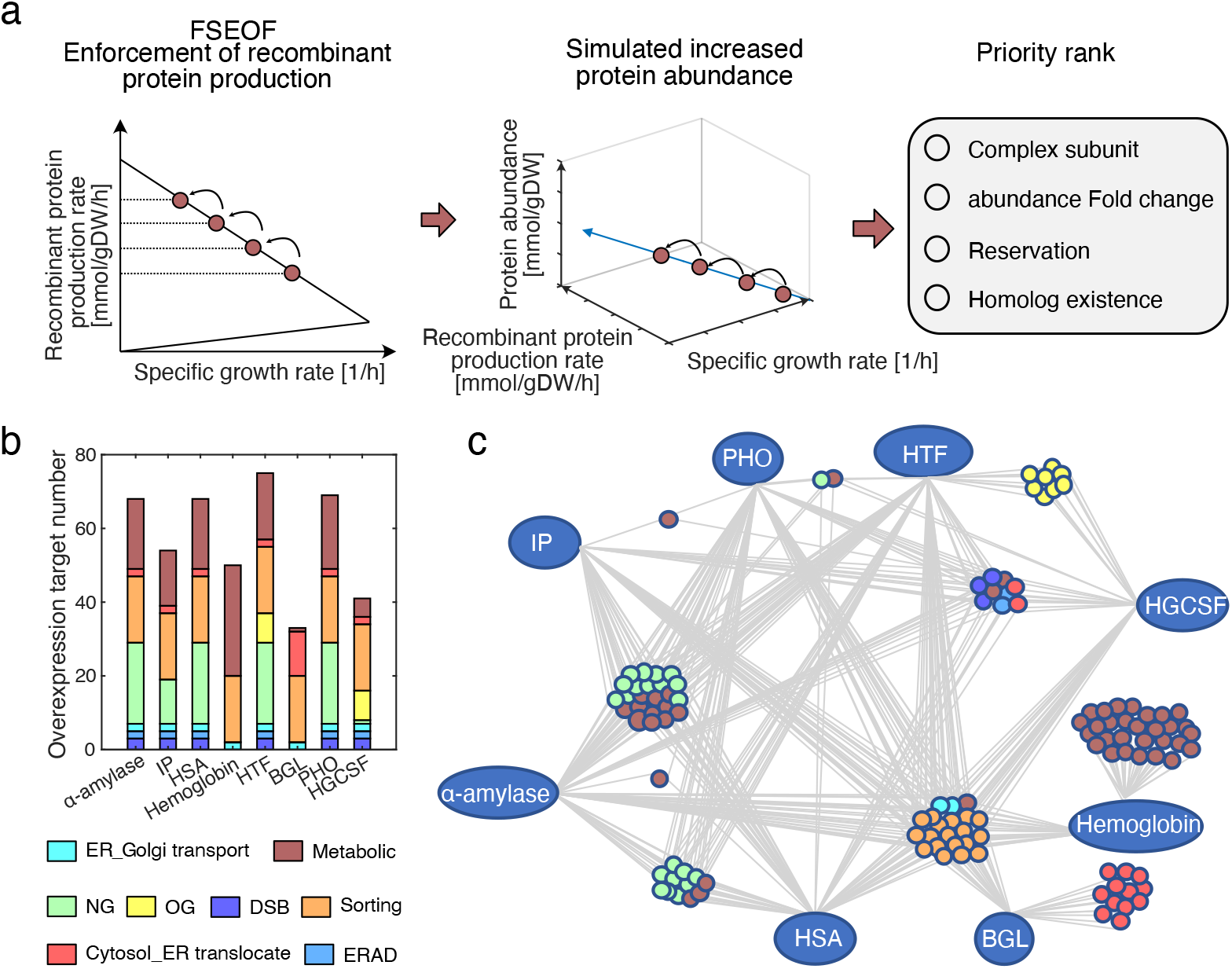
Prediction and comparison of overexpression targets for improving recombinant protein production. a) Adapted FSEOF method for target identification. b) Overview of the predicted overexpression targets for eight recombinant proteins grouped by pathways. c) Comparison of predicted targets for the eight recombinant proteins.

### Protein features impact recombinant protein production

Different secretory proteins utilize different components of the secretory pathway to be processed based on their amino acid composition and PTMs. To identify the factors that influence secreted protein levels, we expanded pcSecYeast to describe the production of eight different recombinant proteins by adding the corresponding recombinant protein production and secretion reactions. These eight recombinant proteins differ in protein size and PTMs (detailed information in Supplementary Table 5). Note that hemoglobin folds with heme as a prosthetic group, which requires balancing of heme biosynthesis and its recombinant protein production (Fig. 4a)^28^. We generated eight specific models to simulate the maximum recombinant protein secretion under various growth rates. We observed that the maximum production rates were achieved at medium-specific growth rates for all the studied recombinant proteins (Fig. 4b), consistent with previous reports of bell shape kinetics for recombinant protein production in *S. cerevisiae* and *Pichia pastoris*^29–31^. Insulin precursor (IP) and α-amylase production were reported as growth dependent^32^, but only for the investigation of a more narrow interval of specific growth rates (0.05-0.2 h^-1^), which is consistent with the model simulations. At high specific growth rates, there is a clear drop of production rate for all recombinant proteins (Fig. 4b), which clearly shows that at high specific growth rates the cell gives priority of its limited capacity of the secretory pathway to native proteins. It is important to note that a default GEM can only describe a linear negative correlation of recombinant protein production with increasing specific growth rates (Supplementary Figure 6). Furthermore, the fact that the simulated α-amylase production by the default GEM is around 1,000 times higher than the experimental values^33^ even with the measured glucose uptake rate as the constraint highlights the huge gaps in default GEM for recombinant protein simulation (Supplementary Figure 6).

**Fig. 6:**
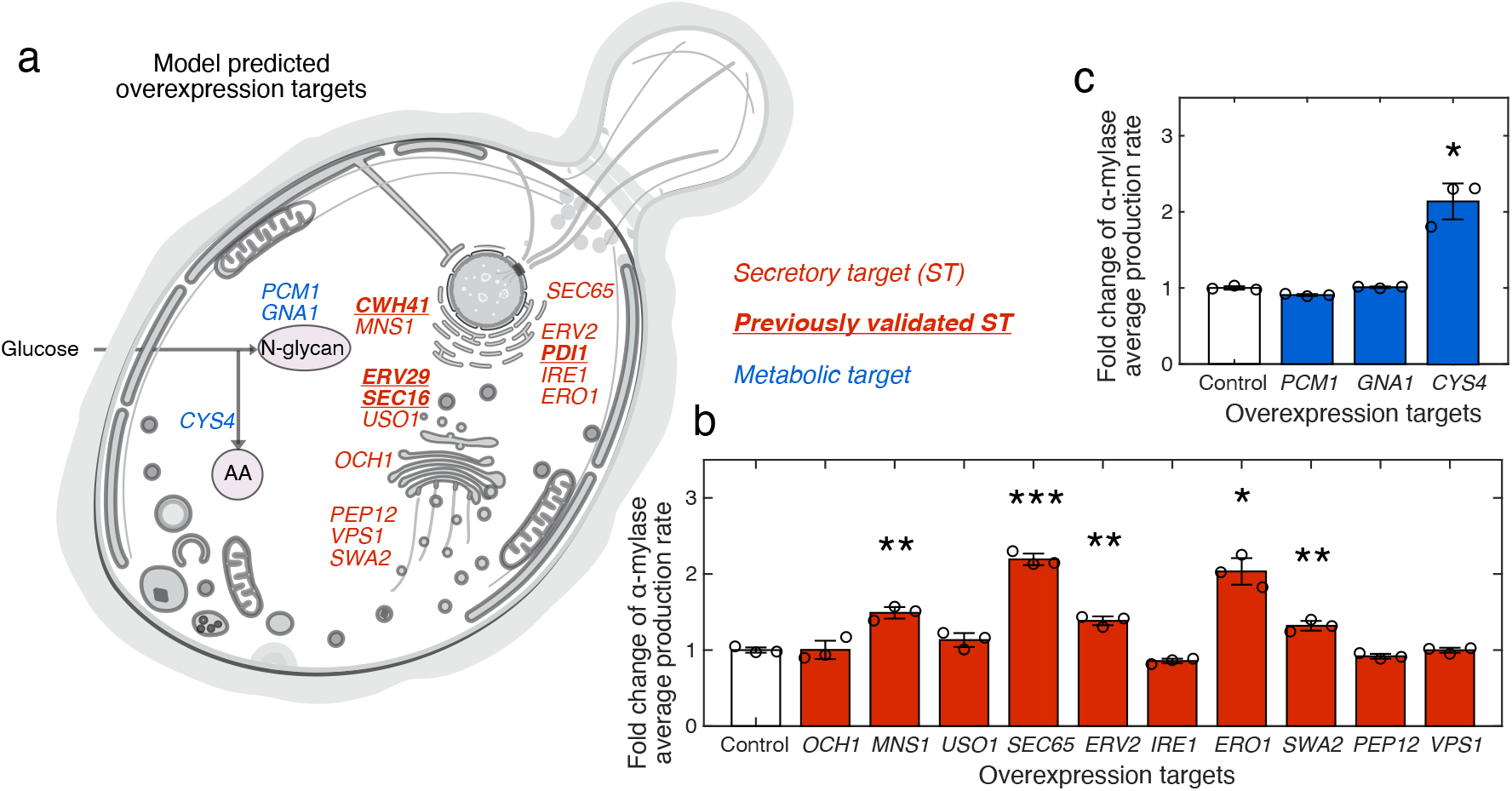
Validation of predicted overexpression targets for α-amylase overproduction. a) Protein localization of the predicted overexpression targets. Yeast compartmentalized figure source: SwissBioPics. b) Validation result of predicted secretory targets. c) Validation result of predicted metabolic targets. *: *P* < 0.05, **: *P* < 0.01, ***: *P* < 0.001. *GNA1* (Glucosamine-6-phosphate acetyltransferase); *PCM1* (PhosphoaCetylglucosamine mutase); *CYS4* (Cystathionine beta-synthase); *CWH41* (Processing alpha glucosidase I); *OCH1* (Mannosyltransferase of the cis-Golgi apparatus); *MNS1* (Alpha-1,2-mannosidase); USO1(Intracellular protein transport protein from ER to Golgi); *SEC65* (Signal recognition particle subunit); *ERV2* (FAD-linked sulfhydryl oxidase); *IRE1* (Serine/threonine-protein kinase/endoribonuclease); *ERO1* (Endoplasmic oxidoreductin-1); *SWA2* (Auxilin-like clathrin uncoating factor); *VPS1* (Vacuolar protein sorting-associated protein); *ERV29* (ER-derived vesicles protein); *PEP12* (Syntaxin); *PDI1* (Protein disulfide-isomerase); *SEC16* (COPII coat assembly protein).

Furthermore, we investigated which protein feature influences recombinant protein production the most. We found that PTMs have an average higher impact on recombinant protein production compared with amino acid composition (Fig. 4c, Supplementary Table 6). Among all simulated features, O-glycosylation and N-glycosylation have larger negative impacts on recombinant protein production, which suggests that having more glycosylation sites would cause more burden for the cell (Fig. 4c).

### FSEOF identifies overexpression targets for recombinant protein overproduction

Identifying engineering targets is crucial to improve the specific recombinant protein production rate. Predicting gene overexpression targets is more difficult and complex than predicting gene deletion targets since amplification of gene expression does not always increase the metabolic fluxes^34^. To fully validate the predictive power of pcSecYeast, we used the generated recombinant protein-specific models to predict overexpression targets for increasing the recombinant protein production. Target prediction was performed using adapted Flux Scanning based on Enforced Objective Function (FSEOF)^34^, where the model was constrained with a stepwise decrease in the specific growth rate, and recombinant protein production was maximized. The original FSEOF method selects fluxes that increase with the enforcement of recombinant protein production in the GEM simulations and identifies those reactions and associated genes as overexpression targets. Since we can compute the protein levels from the pcSecYeast simulations, we can directly select proteins, as overexpression targets, that having increased levels result in increased recombinant protein production (Fig. 5a & Supplementary Dataset). The predicted overexpression targets were ranked for priority and compared among the eight recombinant proteins (Fig. 5b&c). We predicted around 70 overexpression targets for each of the eight recombinant proteins with the majority of them (70%) being in the secretory pathway and 30% in the metabolic part of the model (Fig. 5b&c). Those targets are more likely shared by recombinant proteins when they have the same PTMs. For example, targets in the O-glycosylation pathway are shared by O-glycosylated human-transferrin (HTF) and human granulocyte colony stimulating factor (hGCSF) (Fig. 5c). Surprisingly, even though insulin precursor (IP) contains no N-glycosylation site, some predicted overexpression targets are related to N-glycosylation. This is explained by the fact that N-glycosylation is required for some secretory machinery proteins such as Pdi1 which catalyzes disulfide bond formation in IP production. By removing the disulfide bonds in IP, we found that those N-glycosylation related genes were not predicted as targets (Supplementary Dataset). There are 21 predicted targets shared by all the eight proteins, which are mainly involved in sorting and ER-Golgi transport, suggesting the general importance of these processes in protein secretion (Fig. 5c). We also showed that hemoglobin is the only recombinant protein with multiple unique targets in metabolism, especially for heme production, which demonstrates that metabolism is equally important along with the secretory pathway for improving hemoglobin production. For all the other recombinant proteins, the secretory pathway is more limiting according to the prediction.

### Experimental validation for predicted α-amylase targets

We next validated the predicted overexpression targets for improving α-amylase production. We divided the predicted targets for α-amylase into different groups by their functions and chose 17 targets to validate from all subsystems. There were 14 targets in the secretory pathway spanned in translocation, folding, protein quality control, and sorting subsystems, and three targets in the metabolic part of the model, which are related to N-glycan synthesis and amino acid synthesis (Fig. 6a).

We next sought to test if overexpression of the predicted secretory targets individually could improve the α-amylase production rate. Among them, the glucosidase Cwh41^20^, COPII-coated vesicles proteins Erv29^35^, Sec16^36^ and protein disulfide isomerase Pdi1^35,37^ have already been validated, i.e., overexpression of these proteins can improve α-amylase production and secretion.

As for the remaining ten secretory targets, we performed individual gene overexpression experiments for validation, and found that individual overexpression of *SEC65, MNS1, SWA2, ERV2* and *ERO1* significantly increase the α-amylase production rates by different levels (1.32 to 2.2-fold) (Fig. 6b, Supplementary Table 7). Sec65 is one out of six subunits of the signal recognition particle (SRP), which is involved in protein targeting to the ER^38^. Overexpression of *SEC65* would be anticipated to increase the SRP-dependent co-translational translocation, which would benefit α-amylase translocation from cytosol to ER. Mns1 is involved in folding and ERAD, which is responsible for the removal of one mannose residue from a glycosylated protein. α-amylase contains multiple N-glycosylation sites, and therefore would be benefited from *MNS1* overexpression from facilitated proper folding. *ERO1* encodes a thiol oxidase required for oxidative protein folding in the ER and provides Pdi1 with oxidizing equivalents for disulfide bond formation^39^. We observed that overexpression of *ERO1* also has a positive effect on α-amylase production (2-fold). Besides, overexpressing *ERO1* was able to enhance disulfide-bonded human serum albumin (HSA) secretion in *Kluyveromyces lactis*^40^ and single-chain T-cell receptors (scTCR) and single-chain antibodies (scFv) secretion in *S. cerevisiae*^41^. Therefore, *ERO1* might be considered as a generic target for secretory protein production. *SWA2* is important for vacuole sorting, here we also show that by overexpressing this protein, there is enhancement towards α-amylase production rate (Fig. 6b).

From three metabolic gene targets, only overexpression of *CYS4* led to a significant increase (2.14-fold) of α-amylase productivity (Fig. 6c). Cys4 (Cystathionine beta-synthase) is involved in cysteine synthesis. Comparing the amino acid composition of α-amylase with the average amino acid composition of *S. cerevisiae*, we identified that there is a 9-fold enrichment for cysteine in α-amylase than in the yeast proteome in general (Supplementary Table 8), which explains why overexpression of *CYS4* drastically increases the α-amylase production rate. The other two metabolic targets are Gna1 (Glucosamine-6-phosphate acetyltransferase) and Pcm1 (PhosphoaCetylglucosamine mutase), which are related to the synthesis of N-glycosylation precursor N-linked oligosaccharides. Overexpression of those two genes does not have a significant increase in the α-amylase production rates, which suggests that N-glycosylation precursor synthesis may not be the bottleneck for α-amylase production.

In total, for all the chosen metabolic targets, 1/3 were validated as positive targets, while for identified targets in the secretory pathway, the accuracy was 9/14. Besides the higher accuracy in the secretory targets compared with metabolic targets, FSEOF gives more targets in the secretory pathway even though the fraction of metabolic enzymes in the model is much more than the secretory component. This may give us a hint that for recombinant protein secretion, the secretory pathway is more likely to be the bottleneck, and these results also demonstrate the value of the presented mathematical model for dissecting and systematic analysis of the role of complex protein secretory pathway in recombinant protein production and strain development.

## Discussion

In this study, we presented a genome-scale model of yeast that integrates metabolism, protein translation, protein post-translational-modification, ERAD and sorting processes. The model enables the calculation of ‘unit secretory cost’ for any protein that is processed by the secretory pathway. We have shown that the model can correctly predict the switch from the use of high-affinity to low-affinity glucose transporter as a result of resource optimization (Fig. 2). With the secretory cost calculation and reported transcriptome data, we also detected that upon expression of a recombinant protein which is processed by secretory pathway, yeast optimizes the limited secretory capacity by down-regulating expression of secretory proteins that are expensive to process (Supplementary Figure 2). These two simulations suggest that the cell allocates its limited resources by an optimization strategy, which can be accomplished through regulatory networks that have been tuned through the long evolutionary of yeast upon extracellular and intracellular environments^42–45^.

We next used the model to simulate protein misfolding and retention of CPY and hereby identified that there is a certain ER tolerance to the misfolded protein (Fig. 3). Parameter sensitivity analysis showed the importance of retro-translocation in ER stress. This suggests that increasing the level of retro-translocation may alleviate the ER stress caused by the retention of misfolded protein. Since quality control and ERAD pathways are highly conserved between yeast and higher eukaryotes, this may indicate targets for treating a number of human diseases related to misfolded protein accumulation such as Alzheimer’s and Parkinson’s^46–48^, which has been recently reported as therapeutic interventions^49,50^.

Rational design for recombinant protein production is a crucial task due to the importance of recombinant protein market share and importance, but a very difficult task due to the complexity of the secretory pathway. pcSecYeast serves as a platform for the rational design of system-level engineering targets for recombinant protein production (Fig. 5 & Fig. 6). Besides the experimentally validated the predicted engineering targets for the production of α-amylase (Fig. 6), we also noticed the consistence of the predicted targets for other recombinant proteins with literature reports, such as Hem2, Hem3 and Hem12 for hemoglobin production^28,51^. We also confirmed that even though Hem4 is also in the heme synthesis pathway, this is not a rate-limiting step in the heme synthesis^51^. According to the priority rank from the model prediction, Hem4 has lower priority compared with other proteins such as Hem2 and Hem3. In addition, for targets that were predicted with non-significant impact when overexpressed, we found previous studies to report similar results. For example, overexpressing vacuolar sorting protein Sec15 and Sec4 has been shown to have no positive impact on α-amylase production^36^ (Supplementary Dataset).

To be noted here, our model captures most of the secretory processes, but currently exclude some processes such as Endosome and Golgi-associated degradation pathway (EGAD)^52^, the unfolded protein response and other signaling and regulatory networks^53^. Therefore, including those processes could potentially increase the prediction accuracy in particular when it comes to the dynamic aspects of protein secretion. Besides, we simplified some processes to perform the simulation, which would also introduce some uncertainties, for example, different types of glycans and glycoforms can exist for N-glycosylation^54^. However, modifications to incorporate these processes in the model will be relatively easy in case there is a need to study specific proteins where these processes are important.

In conclusion, we present pcSecYeast as a first genome-scale model which allows systematic modeling of the protein secretory pathway and its interaction with metabolism and gene expression in yeast. This model enables the first time to identify engineering targets for recombinant protein production that can be validated experimentally, and it helps to test the various hypothesis *in silico* for specific protein overexpression. With this new advancement, we expect that this kind of powerful genome-scale secretory model could also be developed for other recombinant protein producing cells, which will entail a fully *in silico* hypothesis generation and identification of cell engineering targets for strain development.

## Methods and materials

### Construction of pcSecYeast and constraint-based analysis

We reconstructed pcSecYeast, which accounts for cell metabolism and protein synthesis processes. Detailed instruction can be found in Supplementary Methods. The reconstruction is based on the latest yeast GEM, Yeast8.3.5^15^. Firstly, we refined all protein PTM precursors synthesis/secretion reactions in the model, such as dolichol synthesis for N-glycosylation, GPI anchor synthesis for GPI modification (Supplementary Table 1). Missing reactions in those precursor synthesis pathways with corresponding GPRs and necessary transport reactions were added into the model for gap-filling.

We split all reversible enzymatic reactions into forward and reverse reactions, and also split reactions catalyzed by isozymes into multiple identical reactions with various isozymes. Besides that, we formulated protein synthesis reactions for all proteins in the model. To facilitate the reconstruction process, the protein synthesis and secretion were divided into 12 different processes: protein translation, protein translocation, ER N-glycosylation, disulfide bond formation, ER O-glycosylation, GPI anchor transfer, COPII anterograde transport, COPI retrograde transport, Golgi N-glycosylation, Golgi O-glycosylation, versatile vesicular transport to destination compartment. We formulated these processes into 123 template reactions. Using the template reactions, we formulated protein synthesis reactions for all proteins in the model. Protein-specific information matrix (PSIM) and localization information for all proteins were downloaded from UniPort^55^ and the SGD^56^ database (Supplementary Table 4). To represent abundance of unpresented proteins that go through ER, we added a dummy ER protein in the model which uses the same composition as the biomass protein, and the PTM for dummy ER protein is calculated as the mean protein modification for proteins that go through ER using the protein abundance from PaxDb^18^ and PSIM information. Protein in the biomass was used to represent protein abundance for proteins excluded in the model. The ratio is rescaled from 1 in original GEM Yeast8 to a lower value 0.3, which was estimated based on the fact that all proteins in the model taking up roughly 70% of the total proteome according to the PaxDb database, which accounts for 4.6% unmodeled dummy ER protein. Detailed model construction and constraints coupling can be found in Supplementary Methods.

### Model simulation for growth using glucose concentration as the constraint

Since the specific growth rate is integrated into the coupling constraints, we adopted a binary search method when we simulated growth. For each specific growth rate, we sampled the glucose concentration until the minimal glucose concentration that can sustain the growth was found. The glucose concentration was used to calculate import rate using the Michaelis– Menten equation where *K*_M_ and maximal *k*_cat_ of glucose transporters were collected from the literature^57–59^. As for the glucose transporters which does not have any *k*_cat_ values, the V_max_ data was used to convert to *k*_cat_ values with the assumption that the expression levels are comparable in the collected dataset since they expressed transporter constructs under constitutive promoters in a yeast glucose-transporter null-mutant^58,60,61^. The model was set with minimal media and the dummy protein production was set as the objective. Besides all mentioned basic constraints in the Supplementary Methods, we added constraints on the fraction of ER membrane proteins and ER volume to avoid the possibility of an unrealistic ER volume. Due to the requirement of the linear programming (LP) solver (SoPlex), all constraints were written in a LP file for solving in each simulation. This method for adding constraints is used in all following simulations unless otherwise stated.

### Estimation of ‘unit secretory cost’ and ‘direct cost’ for secretory proteins

‘Unit secretory cost’ of synthesizing about ∼500 proteins that localize to the cell membrane or are secreted were estimated using the model. At a specific growth rate of 0.1 h^-1^, we used pcSecYeast to produce a sequential small fraction production of those proteins, respectively. The glucose uptake rate minimization was set as the objective. Using the simulated glucose uptake rates and the production rates, we could fit the linear equation to get the slope which is the ‘unit secretory cost’ for each protein. This cost stands for the energetic cost for synthesizing the protein, PTM, sorting and even the related cost for the corresponding fraction of the catalytic machineries in these processes.

‘Direct cost’ accounts for the energetic cost for synthesizing the amino acids, bounded glycan precursors and enzyme bounded energetic molecules, which was calculated with only the default GEM constraint including the mass balance and reaction bound, without any enzyme-related constraint. Since this simulation only require any extra constraint, we used the optimize function and default solver in COBRA toolbox rather than the SoPlex and LP file method.

### Analysis of gene expression versus protein ‘unit secretory cost’

Absolute transcriptome data for three strains (AAC, MH34 and B184) with different α-amylase production levels were used for the correlation analysis (Supplementary Table 9)^20^. Pearson correlation coefficient was used to assess the correlation of ‘unit secretory cost’ with the expression levels.

### Simulation of protein misfolding and accumulation

We used CPY as an example to show how the model responds towards misfolded protein production. CPY was expressed in the model with different levels from the native abundance towards its 25 fold as reported in the literature^26^ by constraining its translation flux. In order to identify the factor causing the accumulation of misfolded protein in ER, we performed the parameter sensitivity analysis for ERAD capacity, ER volume, ER membrane space, total secretory machinery capacity and retro-translocation enzyme abundance, respectively. Since the membrane space and the volume of proteins are positively correlated with the protein weight^62^, ER membrane space and ER volume constraints can be converted to proteome abundance constraints, which can be calculated from the proteome data. Therefore, all these parameters can be constrained by an upper limit on the total abundance of the corresponding proteins. In the meanwhile, we changed the misfolding ratio constraint of CPY by coupling the flux of misfolding reaction and the translation reaction of CPY. When misfolded protein was retained in the ER, we used the multiple round reactions of binding Kar2 and Pdi1 to reflect its occupancy of Kar1 and Pdi1 as reported^26,39^. The coefficient of this reaction was used to represent the time for the retention. For the combination of CPY expression levels and misfolding ratio, we used the binary search as mentioned above to search for the maximum specific growth rate. The accumulated CPY rate was obtained from the simulated flux under the found maximum growth rate condition. To reflect the CPY production as close to the *in vivo* as possible, we adjusted the N-glycans attached to the N-glycosylation sites of CPY as reported^63^.

### Expansion of pcSecYeast to recombinant protein specific models

We expanded pcSecYeast to represent the recombinant protein production by adding the production and secretion reactions using the same template reactions for the native proteins. The PTMs, amino acid sequence and leader sequence were collected from the literature. Detailed information for those proteins and the literature reference can be found in Supplementary Table 5.

### Model simulation for recombinant protein production

To simulate recombinant protein production, the model was constrained with a certain specific growth rate, and then the protein production was maximized. SD-2×SCAA medium was used in the simulations^33^. All constraints mentioned were added when writing the LP file for solving by SoPlex.

### Protein feature importance analysis

Machine learning was integrated to score the importance of factors. In this study, various factors (PTMs, amino acid compositions) were used as the input features and the maximum recombinant protein production rate was used as the target label. We split the created dataset into a training dataset and testing dataset at the ratio of 80% and 20%, respectively. A random forest regressor with 10 estimators was used to train the model. Feature importance scores from the random forest were computed by SHAP (SHapley Additive exPlanations)^64^.

### Overexpression target prediction for recombinant protein overproduction

Identification of overexpression targets for improving recombinant protein production was performed using the concept of FSEOF^34^ but to identify the proteins with increased expression during the enforcement of recombinant protein production. To be noted here, original FSEOF searches for the candidate fluxes to be amplified through scanning for those fluxes that increase with enforced product formation flux under the objective function of maximizing biomass formation flux, which is under the assumption that there is a tradeoff between growth and target production. pcSecYeast is much more complex than the default GEM and can better represent the cell state which the recombinant protein production does not always increase with the decrease of growth. Besides that, there is metabolic state switch of the fermentation ratio for energy production. Therefore, to eliminate growth and metabolic state influence, we selected a small window (0.25h^-1^-0.3 h^-1^) for this analysis. At each growth rate in this window, we maximized the recombinant protein production rate without any constraint on exchange rates. Proteins with amplificated expression accompanied increased recombinant protein production were selected as initial overexpression targets. Then, we used several cutoffs to rank the targets further: 1) for proteins that always increase with the enforcement of the recombinant protein production with a Pearson correlation score over 0.9, the priority score was set to 1; 2) for proteins with priority score 1 and showed 1.3-fold change of the maximum recombinant protein production state towards the maximum specific growth rate, the priority score was set to 2; 3) for proteins with priority score 2 and showed a comparable difference towards the reference PaxDb abundance, which represents the reservation state of the protein abundance in the cell, the priority score was set to 3; 4) for proteins with priority score 3 and were neither subunits of complexes nor contain homologs, the priority score was set to 4. Targets with higher priority scores should be prioritized. Proteins with the priority score 0 in the result indicate those proteins are not identified as overexpression targets. Based on the criteria, we ranked the targets and generated an annotated table as result for all eight recombinant proteins (Supplementary Dataset). For plotting the common targets shared by all eight recombinant proteins analyzed in this study, we only chose the priority score of 3 and 4 for the analysis.

## Experimental validation

### Strains and plasmids

All strains and plasmids used in this study are listed in Supplementary Table 10. Plasmids for gene overexpression were constructed by insertion of the gene fragment, which was amplified from the yeast genome then assembled with the expression vector pSPGM1 through Gibson assembly method. The standard LiAc/SS DNA/PEG method was used for yeast transformation.

### Media and culture conditions

For strain constructions, yeast strains were grown in SD-URA medium at 30 °C according to the auxotrophy of the cells. For α-amylase production in shake flasks, yeast strains were cultured for 96 h at 200 rpm, 30 °C with an initial OD_600_ of 0.05 in the SD-2×SCAA medium containing 20 g/l glucose, 6.9 g/l yeast nitrogen base without amino acids, 190 mg/l Arg, 400 mg/l Asp, 1,260 mg/l Glu, 130 mg/l Gly, 140 mg/l His, 290 mg/l Ile, 400 mg/l Leu, 440 mg/l Lys, 108 mg/l Met, 200 mg/l Phe, 220 mg/l Thr, 40 mg/l Trp, 52 mg/l Tyr, 380 mg/l Val, 1 g/l BSA, 5.4 g/l Na_2_HPO_4_ and 8.56 g/l NaH_2_PO_4_.H_2_O (pH=6.0)^33^.

### α-Amylase quantification

The α-amylase activity was measured using the α-amylase assay kit (Megazyme) with a commercial α-amylase from *Aspergillus oryzae* (Sigma-Aldrich) as the standard. Samples were centrifuged for 10 min at 15,000 g, 4 °C and the supernatant was used for extracellular α-amylase quantification.

## Code availability

To facilitate further usage, we provide all codes and detailed instruction in GitHub repository: https://github.com/SysBioChalmers/pcSecYeast. All codes to reproduce figures were also included in the GitHub repository.

## Data availability

All data used in this study are included in supplementary files and GitHub repository.

## Author contribution

F.L. and J.N. designed the research. F.L. performed the research. Y.C. contributed to the model simulation. Q.Q. and Y.W. performed the experimental validation. L.Y. contributed to the protein feature importance analysis. I.EE. contributed to the model reconstruction. F.L., Y.C., Q.Q., Y.W., A.F., I.E., E.JK. and J.N. analyzed the data. F.L., Y.C., E.JK. and J.N. wrote the paper. All authors approved the final paper.

## Acknowledgement

This project has received funding from the Novo Nordisk Foundation (grant no. NNF10CC1016517), VINNOVA center CellNova (2017-02105), the Knut and Alice Wallenberg Foundation, and the European Union’s Horizon 2020 research and innovation program with projects DD-DeCaF (grant no. 686070). The computations were enabled by resources provided by the Swedish National Infrastructure for Computing (SNIC) at Chalmers Centre for Computational Science and Engineering (C3SE) and High Performance Computing Center North (HPC2N), partially funded by the Swedish Research Council through grant agreement no. 2018-05973.

