## Supplementary Figure for "Genome scale modeling of the protein secretory pathway reveals novel targets for improved recombinant protein production in yeast"

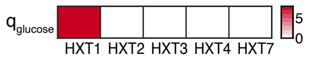


Supplementary Figure 1: Fluxes (mmol/gCDW/h) carried by glucose transporters at maximum growth simulation when both Hxt1 and Hxt7 are set with the $k_{\mathrm{cat}}$ value for Hxt7 (197/s).

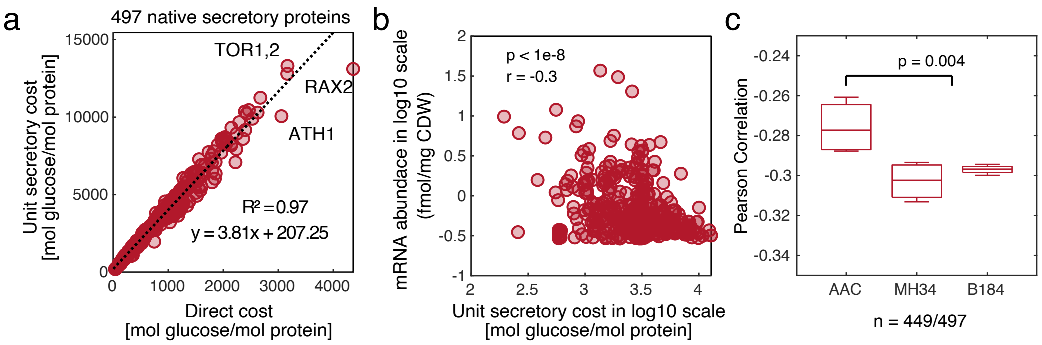


Supplementary Figure 2: *S. cerevisiae* suppress costly native proteins upon recombinant protein production. a) Simulated secretory costs for 497 native secretory proteins in *S. cerevisiae*. Direct cost includes the energetic cost for synthesis, modification and secretion of this protein, while unit secretory cost additionally includes the cost for the corresponding increased fraction of the catalytic machineries in these processes caused by the increase of this protein. b) Example of negative Pearson correlation of mRNA levels with 'unit secretory costs' of native secretory and cell membrane proteins in AAC strain at $0.1h^{-1}$. c) Pearson correlation of mRNA level with glucose cost of native secretory proteins for three α-amylase strains. *P* value for the AAC with the MH34 and B184 was calculated using Wilcoxon rank sum test. AAC: low yield α-amylase strain; MH34 and B184: high yield α-amylase strain. 449 out of 497 native proteins have measured mRNA abundances and were used in the correlation analysis. Each point in the box plot represents one correlation and has *P* value < 1e-8.

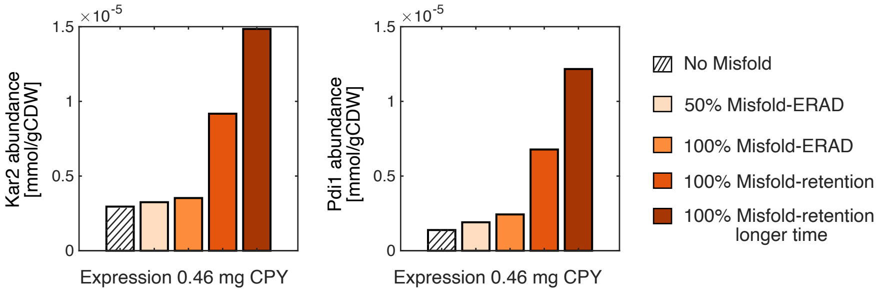


Supplementary Figure 3: Simulated abundances of Kar2 and Pdi1 in expression of 0.46 mg CPY with different routes.

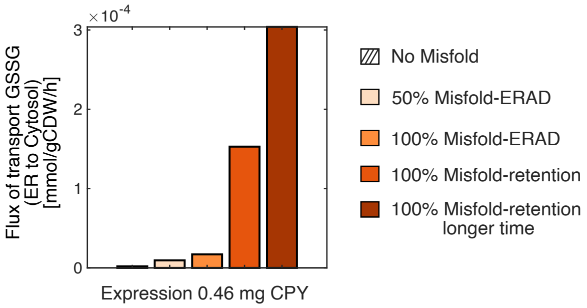


Supplementary Figure 4: Simulated fluxes for transporting GSSG from ER to cytosol in expression of 0.46 mg CPY with different routes.

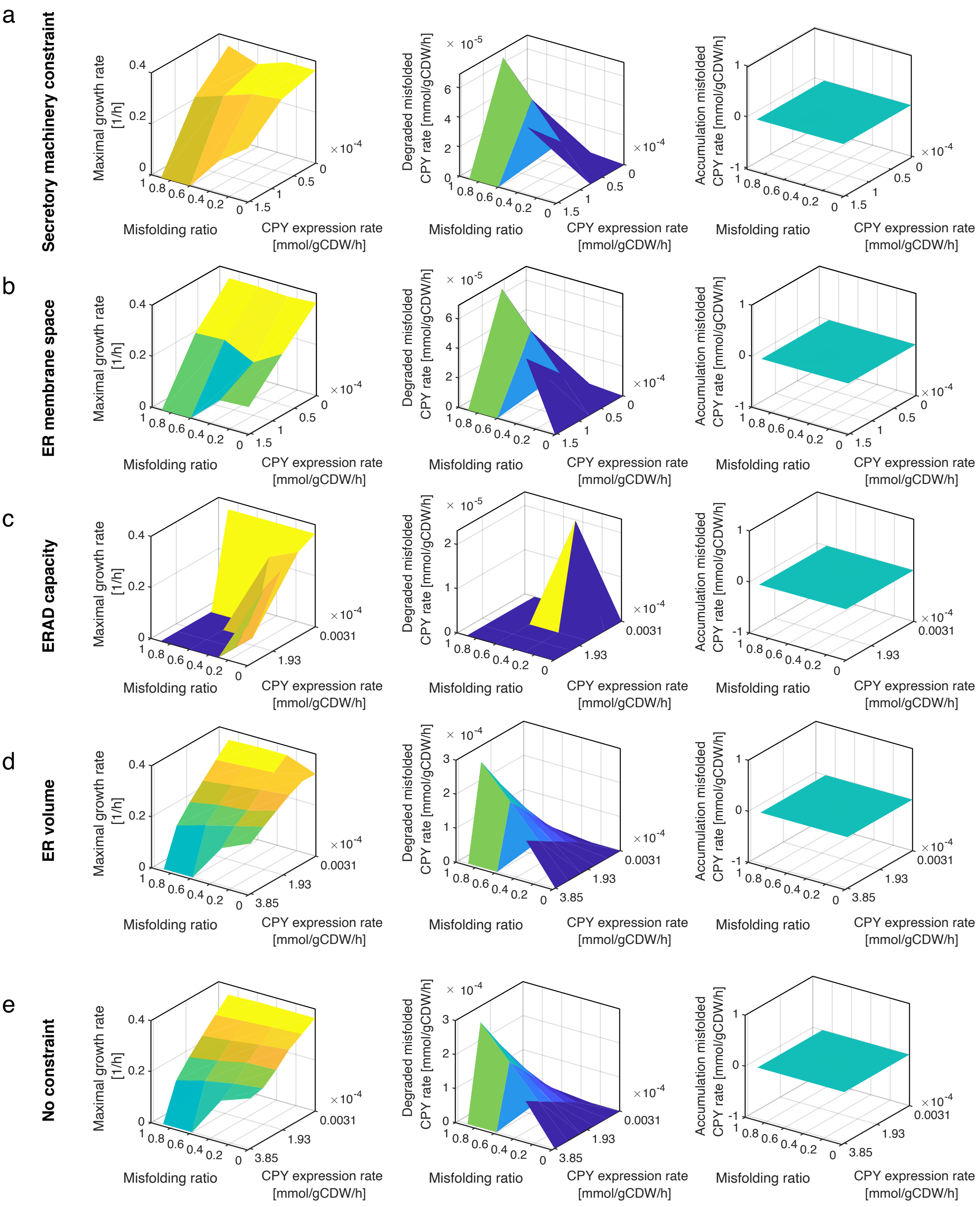


Supplementary Figure 5: Parameter analysis for the accumulation of misfolded CPY.

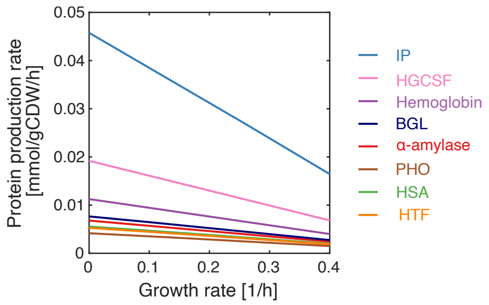


Supplementary Figure 6: Simulation of recombinant protein production under diverse specific growth rates using Yeast8 expanded with a reaction for production of the recombinant protein.
