## Supplementary Methods for "Genome scale modeling of the protein secretory pathway reveals novel targets for improved recombinant protein production in yeast"

### Model Generation and Formulation

### Summary

This manual describes how the data were collected, how the model was reconstructed, and how the optimization problem was generated. In the beginning, we write an index for the main Excel and MATLAB files used to generate and simulate the model. All excel files can be found in the Github repository [https://github.com/SysBioChalmers/pcSecYeast].

#### Table S1.xlsx

contains all collected information.

- **Annotation & Annotation_extra:** gene names and sequence information of all *S. cerevisiae* proteins.
- **Machinery:** proteins which makes ribosome, ribosome assembly complex and proteasome complex
- **Secretory:** proteins which makes secretory machinery complexes
- **Secretory_ref:** references for how those secretory complexes are summarized
- **kdeg:** kdeg information collected from reference[1]
- **kcat_info_metabolic:** $k_{cat}$ values used in the metabolic part
- **kcat_info_sectory:** $k_{cat}$ values used in the secretory part
- **kcat_info_machinery:** $k_{cat}$ values used in the ribosome, ribosome assembly and proteasome

#### Protein_information.xlsx

contains the Protein Specific Information Matrix (PSIM) info for all proteins in *S. cerevisiae*, including information such as whether the protein go through ER, existence of the signal peptide, number of disulfide bond sites, N-glycosylation sites, O-glycosylation sites, transmembrane domains, GPI sites, protein localization, protein amino acid sequence and length, signal peptide sequence and length. Information were collected from literature[2] and UniPort database[3],

#### Main MATLAB file:

Run the function `buildModel.m` to get pcSecYeast model.

### 1. Information collection

#### 1.1 Protein sequence

We downloaded protein sequence information from the UniProt database[3], which is stored as in the Annotation&Annotation_extra sheet in Table S1. This information is stored as a mat file in `Protein_sequence.mat` in the GitHub repository.

#### 1.2 Protein Information

All protein information was collected from the literature[2] and UniProt database[3]. For proteins which exist in multiple compartments, we chose the first annotated compartment as its localization. As for proteins which denoted have signal peptide but does not annotate the signal peptide length, we used the first 21 aa as its signal peptide.

#### 1.3 Protein stoichiometry

For each functional protein, we should determine whether its functional unit is a monomer or oligomer. To do so, we collected protein stoichiometry information from PDBe database (https://www.ebi.ac.uk/pdbe/) as well as Complex Portal website (https://www.ebi.ac.uk/complexportal/home). This information is stored as a mat file in `Protein_stoichiometry.mat` in the GitHub repository.

### 2. reformulation of metabolic model

#### 2.1 Metabolic model origin

The latest GEM Yeast8[4] for *S. cerevisiae* was used as the basis for the model pcSecYeast. Yeast8.3.5 was extracted from the GitHub: https://github.com/SysBioChalmers/yeast-GEM.

#### 2.2 Adding reactions for precursor production of post-translational modification

Yeast8.3.5 was firstly curated by adding reactions to ensure the production of all precursors for the secretory pathway such as glycans. Those reactions can be found in the excel `Yeast8_Modification.xlsx` in the GitHub repository. In total 92 reactions were added into Yeast8.3.5, which contains glycan synthesis, GPI anchor synthesis and transport reactions to shuttle currency metabolites between cytosol and other compartments in the secretory pathway. As for the GPI anchor synthesis, we chose the 1-phosphatidyl-1D-myo-inositol (1-16:0, 2-18:1) as the starting phosphatidyl-inositol according to the literature report[5]. Other lipids attached to GPI anchor was according to the reference[6]. Function `modifyYeast8` was used to add these reactions into Yeast8.

##### 2.3 Change reversibility

The updated model was reformulated by splitting reversible reactions into forward and reverse reactions and splitting reactions catalyzed by isozymes into multiple identical reactions with various isozymes. This step was performed to facilitate later enzyme constraining step. Function `splitModel` was used to perform this change.

#### 3. Protein related process reconstruction

##### 3.1 Peptide translation

We formulated the translation process for all proteins in the model. The substrates of translation include charged amino acid tRNAs, while the products include uncharged tRNA and the translated peptide. Besides the charged tRNAs, the translation process requires energy. Three steps in protein translation were considered: translation initiation, translation elongation and translation termination. The translation initiation process requires one ATP molecule for binding mRNA with initiation factors, one ATP molecule for every step in the scanning process, and two GTP molecules for initiation factors. The translation elongation process has been studied intensely in yeast[7]. In summary, for each amino acid, two ATPs and two GTPs are required during the elongation. As for the energy cost for the translation termination, one ATP and one GTP are required for each peptide. The energy cost is calculated as 2N+3 ATP and 2N+3 GTP for each peptide, where N is the number of amino acids for that peptide. The energy molecules were included as substrates in the translation reactions. Note that we simplified the model by assuming that all proteins are translated in the cytoplasm by the ribosome. Since the charged and uncharged tRNA are already metabolites in the original Yeast8, we did not update the tRNA charging process. Function `addTranslationRxns` is used to add translation reactions for proteins.

Example of translation reaction for protein YAL012W.

reaction id: r_YAL012W_peptide_translation

reaction equation: 1582 H2O[c] + 791 ATP[c] + 791 GTP[c] + 41 Ala-tRNA(Ala)[c] + 13 Arg-tRNA(Arg)[c] + 22 Asn-tRNA(Asn)[c] + 21 Asp-tRNA(Asp)[c] + 15 Gln-tRNA(Gln)[c] + 21 Glu-tRNA(Glu)[c] + 28 Gly-tRNA(Gly)[c] + 14 His-tRNA(His)[c] + 24 Ile-tRNA(Ile)[c] + 40 Leu-tRNA(Leu)[c] + 22 Lys-tRNA(Lys)[c] + 4 Met-tRNA(Met)[c] + 14 Phe-tRNA(Phe)[c] + 17 Pro-tRNA(Pro)[c] + 29 Ser-tRNA(Ser)[c] + 28 Thr-tRNA(Thr)[c] + 2 Trp-tRNA(Trp)[c] + 10 Tyr-tRNA(Tyr)[c] + 30 Val-tRNA(Val)[c] -> 1582 H+[c] + 1582 phosphate[c] + 791 ADP[c] + 791 GDP[c] + 41 tRNA(Ala)[c] + 13 tRNA(Arg)[c] + 22 tRNA(Asn)[c] + 21 tRNA(Asp)[c] + 15 tRNA(Gln)[c] + 21 tRNA(Glu)[c] + 28 tRNA(Gly)[c] + 14 tRNA(His)[c] + 24 tRNA(Ile)[c] + 40 tRNA(Leu)[c] + 22 tRNA(Lys)[c] + 4 tRNA(Met)[c] + 14 tRNA(Phe)[c] + 17 tRNA(Pro)[c] + 29 tRNA(Ser)[c] + 28 tRNA(Thr)[c] + 2 tRNA(Trp)[c] + 10 tRNA(Tyr)[c] + 30 tRNA(Val)[c] + YAL012W_peptide[c]

catalyst: Mach_Ribosome_complex

#### 3.2 Protein processing and complex formation for proteins that do not go through the secretory pathway

Translated nascent peptides need to be folded to form functional complexes for performing certain functions. As for proteins that do not go through the secretory pathway, we added a simplified process for folding, misfolding and complex formation, similar to what has been done in proteome-constrained models for *S. cerevisiae*[8] and *Lactococcus lactis*[9]. The folding process in the model happens in the cytosol and complex formation happens in the localization of the complex. As for complexes with multiple subunits, the stoichiometry information collected before was used in the complex formation reaction.

Example of protein processing and complex formation for YBR153W

reaction id: YBR153W_folding_c

reaction equation: YBR153W_peptide[c] -> YBR153W_folding[c]

catalyst: -

```
reaction id: YBR153W_degradation_misfolding_c
reaction equation: YBR153W_misfolding[c] -> YBR153W_subunit[c]
catalyst: -

reaction id: YBR153W_misfold_c
reaction equation: YBR153W_folding[c] -> YBR153W_misfolding[c]
catalyst: -

reaction id: YBR153W_dilution_misfolding_c
reaction equation: YBR153W_misfolding[c] ->
catalyst: -

reaction id: r_0015_complex_formation
reaction equation: 2 YBR153W_folding[c] -> r_0015_complex[c]
catalyst: -
```

#### 3.3 Protein processing and complex formation for proteins that go through the secretory pathway

As for proteins that go through the secretory pathway, folding and further processing were detailly described in the model.

##### 3.3.1 Protein translocation

The nascent translated peptide would be translocated into the ER for further modification. There are three different pathways for protein translocation from the cytosol to the ER: co-translational translocation, post-translational translocation, and post-translational translocation-tail-targeting[10,11].

Co-translational translocation in *S. cerevisiae* involves the interplay of the nascent peptide, the ribosome, the signal recognition particle (SRP), the signal recognition receptor (SR) and either the Ssh1 or the Sec61 translocon pore. SRP is a complex of six proteins (Srp14, Srp21, Srp54, Srp68,

Srp72, and Sec65) and a 7S single RNA (SCR1)[12–14]. SRP interacts with the signal peptide of nascent peptides to form ribosome–nascent chain (RNC) complex, thus by the interaction of SPR and a signal receptor complex (SR), encoded by *SRP101* and *SRP102,* the RNC complex can attach to ER membrane. Finally, the RNC complex is transferred to the translocon, and then SRP and SR dissociate. Nascent peptides are then translocated into the ER. GTP bind to both SRP (via the Srp54 subunit) and the SR, which is critical for their interaction.

In the model, this process was divided into six template reactions. Only the protein with signal peptide adopted this pathway, and if the signal peptide sequence has not been annotated, we use the first 21 amino acids of the protein as the signal peptide.

1. signal peptide recognition
2. ER receptor biding to peptide-SRPC
3. binding of peptide -SRPC-SRC to the translocator (SEC61C)
4. binding of peptide -SRPC-SRC to the translocator (SSH1C)
5. signal peptidase
6. export the signal peptide out of ER for degradation

Example of co-translation reaction of YDR367W

reaction id: YDR367W_co_translation_TC_sec_SRPC_complex
reaction equation: YDR367W_peptide[c] -> YDR367W_tanslocate_1[c]
catalyst: sec_SRPC_complex

reaction id: YDR367W_co_translation_TC_sec_SRC_complex
reaction equation: YDR367W_tanslocate_1[c] -> YDR367W_tanslocate_2[c]
catalyst: sec_SRC_complex

reaction id: YDR367W_co_translation_TC_sec_SEC61C_complex
reaction equation: 2 H2O[c] + 2 GTP[c] + YDR367W_tanslocate_2[c] -> H+[c] + 2 phosphate[c] + 2 GDP[c] + YDR367W_tanslocate_3[c]
catalyst: sec_SEC61C_complex

reaction id: YDR367W_co_translation_TC_sec_SSH1C_complex

reaction equation: 2 H2O[c] + 2 GTP[c] + YDR367W_tanslocate_2[c] -> H+[c] + 2 phosphate[c] + 2 GDP[c] + YDR367W_tanslocate_3[c]

catalyst: sec_SSH1C_complex

reaction id: YDR367W_co_translation_TC_sec_SPC_complex

reaction equation: H2O[c] + YDR367W_tanslocate_3[c] -> YDR367W[er] + YDR367W_sp[er]

catalyst: sec_SPC_complex

reaction id: YDR367W_export_sp_to_c

reaction equation: YDR367W_sp[er] -> YDR367W_sp[c]

catalyst: -

Post-translational translocation is equivalently important with co-translational translocation. Nascent peptides exit the ribosome with the help of RAC chaperones (Ssb1, Ssz1 and Zuo1)[15]. After that, nascent peptides remain in an unfolded or loosely folded state, bind to the cytosolic chaperones Ssa1 and Ydj1 to avoid aggregation, which would be released before the translocation initiate. The translocation is mediated by the SEC complex, which comprises Sec61, Sbh1, Sss1, Sec62, Sec63, Sec71, and Sec72. The chaperon Kar2 in the ER lumen was suggested to drive the nascent protein into the ER[16].

In the model, this process was divided into four template reactions. Coefficient of ATP in step 4 was set as length/40, based on the assumption that an ATP molecule that bound to the chaperone Kar2, is hydrolysed to ADP for every 40 amino acids that pass through the translocon pore[17].

1. exit the ribosome
2. bind to the cytosolic chaperone
3. translocation
4. pulling of nascent protein

Example of post-translation reaction of YDR453C

reaction id: YDR453C_Post_translation_PSTA_sec_RAC_complex
reaction equation: YDR453C_peptide[c] -> YDR453C_tanslocate_1[c]
catalyst: sec_RAC_complex

reaction id: YDR453C_Post_translation_PSTA_sec_Ssa1_Ydj1_Snl1_complex
reaction equation: H2O[c] + ATP[c] + YDR453C_tanslocate_1[c] -> H+[c] + phosphate[c] + ADP[c] + YDR453C_tanslocate_2[c]
catalyst: sec_Ssa1_Ydj1_Snl1_complex

reaction id: YDR453C_Post_translation_PSTA_sec_SEC61SEC63C_complex
reaction equation: YDR453C_tanslocate_2[c] -> YDR453C_tanslocate_3[c]
catalyst: sec_SEC61SEC63C_complex

reaction id: YDR453C_Post_translation_PSTA_sec_BIP_NEFS_complex
reaction equation: 5 H2O[c] + 5 ATP[c] + YDR453C_tanslocate_3[c] -> 5 H+[c] + 5 phosphate[c] + 5 ADP[c] + YDR453C[er]
catalyst: sec_BIP_NEFS_complex

Post-translational translocation-tail targeting is a special translocation process, especially for tail-anchored (TA) proteins[11]. This process is also termed as the GET pathway, which involves Sgt2, Get4, Get5 and Get3 proteins. This process initiates from loading the TA proteins from the ribosome by the complex composed of Sgt2, Get4 and Get5. Then, the complex binds to Get3, a cytosolic transmembrane domains (TMD) recognition complex. This Get3 complex would deliver the protein to the ER receptor composted of Get1 and Get2.

This process was formulated into three template reactions in the model, proteins with GPI anchor adopt this pathway for translocation.

1. load the TA proteins
2. bind to Get3
3. bind to ER receptor

Example of post-translation reaction-tail-targeting of YBL011W

reaction id: YBL011W_Post_translation_PSTB_sec_Sgt2_Get4_Get5_complex
reaction equation: YBL011W_peptide[c] -> YBL011W_tanslocate_1[c]
catalyst: sec_Sgt2_Get4_Get5_complex

reaction id: YBL011W_Post_translation_PSTB_sec_Get3_complex
reaction equation: H2O[c] + ATP[c] + YBL011W_tanslocate_1[c] -> H+[c] + phosphate[c] + ADP[c] + YBL011W_tanslocate_2[c]
catalyst: sec_Get3_complex

reaction id: YBL011W_Post_translation_PSTB_sec_Get1_Get2_complex
reaction equation: YBL011W_tanslocate_2[c] -> YBL011W[er]
catalyst: sec_Get1_Get2_complex

3.3.2 Disulfide bond formation

For proteins annotated with disulfide bonds, reactions of this step were added. The nascent peptide is captured by the chaperone Kar2, which mediates folding. Sulfhydryl groups are then oxidized by protein disulfide isomerases (Pdi1). Reoxidation of Pdi1 is mediated by ER oxidoreductin (Ero1), which in turn transfers electrons to $O_2$, thereby generating reactive oxygen species (ROS)[10].

This step in the model was divided into two template reactions.

1. bind to the chaperone
2. disulfide bond formation

Example of disulfide bond formation of YCL035C

reaction id: YCL035C_DSB_sec_BIP_NEFS_complex
reaction equation: 2.775 H2O[er] + 2.775 ATP[er] + YCL035C[er] -> 2.775 H+[er] + 2.775 phosphate[er] + 2.775 ADP[er] + YCL035C_Kar2ATPcplx[er]
catalyst: sec_BIP_NEFS_complex

reaction id: YCL035C_DSB_PDI_II_sec_PDI1_ERV2_Ero1p_complex
reaction equation: PDI-ox[er] + YCL035C_Kar2ATPcplx[er] -> PDI[er] + YCL035C_DSB[er

catalyst: sec_PDI1_ERV2_Ero1p_complex

3.3.3 GPI formation

GPI formation process was formulated into several reactions according to reference[5]. Metabolic reactions for glycosylphosphatidylinositols (GPIs) synthesis were gap-filled in the Yeast8 modification step. Here, the first step is to transfer synthesized GPIs into proteins, followed by several remodeling steps of the sugar and lipid moieties in the GPI anchor. The acyl chain from the inositol is firstly removed by the Bst1, then a phospholipase A2 (Per1) removes the C18:1 fatty acid of the primary anchor[18]. After that, a C26:0 fatty acid is attached to the diacylglycerol moiety by Gup1. For most GPI anchors, this modified diacylglycerol-based anchor is subsequently transformed into a ceramide-containing anchor by Cwh43[19]. Ted1 further removes a phosphoethanolamine (PEtN) on the second mannose. Even though the GPI anchor will be further modified in the Golgi, we do not include the Golgi modification part as the enzyme responsible for this has not been identified. This could be easily added into the model in the future[20].

In the model, the GPI formation was divided into six steps:

1. GPI transfer
2. removal of the acyl chain from the inositol
3. removal of the unsaturated acyl chain at the sn-2 position of diacylglycerol to form lyso-GPI
4. transfer C26 saturated acyl chain to the sn-2 position
5. change the lipid moiety to ceramide consisting of PHS with a hydroxy-C26 fatty acid
6. removes a PEtN on the second mannose
7. GPI transfer with recycled GPI anchor from misfolded protein

Example of GPI formation and transfer of YDR437W

reaction id: YDR437W_GPIRI_sec_GPIR_complex
reaction equation: 6-O-2-O-((2-aminoethyl)phosphoryl)-alpha-D-mannosyl-(1-2)-{alpha-D-mannosyl-2-O-((2-aminoethyl)phosphoryl)-(1-2)-alpha-D-mannosyl-(1-6)-2-O-((2-aminoethyl)phosphoryl)-alpha-D-mannosyl-(1-4)-alpha-D-glucosaminyl}-O-acyl-1-phosphatidyl-1D-myo-inositol[er] + YDR437W[er] -> H2O[er] + YDR437W_GPI_G1[er]

catalyst: sec_GPIR_complex

reaction id: YDR437W_GPIRII_sec_Bst1p_complex
reaction equation: YDR437W_GPI_G1[er] -> palmitate[erm] + YDR437W_GPI_G2[er]
catalyst: sec_Bst1p_complex

reaction id: YDR437W_GPIRIII_sec_Per1p_complex
reaction equation: H2O[er] + YDR437W_GPI_G2[er] -> oleate[er] + YDR437W_GPI_G3[er]
catalyst: sec_Per1p_complex

reaction id: YDR437W_GPIRIV_sec_Gup1p_complex
reaction equation: hexacosanoyl-CoA[er] + YDR437W_GPI_G3[er] -> H+[er] + coenzyme A[er] + YDR437W_GPI_G4[er] catalyst: sec_Gup1p_complex

reaction id: YDR437W_GPIRV_sec_Cwh43p_Gpi7p_Mcd4p_complex
reaction equation: ceramide-3 (C26)[er] + YDR437W_GPI_G4[er] -> diglyceride (1-26:0, 2-16:0)[er] + YDR437W_GPI_G5[er]
catalyst: sec_Cwh43p_Gpi7p_Mcd4p_complex

reaction id: YDR437W_GPIRVI_sec_Ted1p_complex
reaction equation: H2O[er] + YDR437W_GPI_G5[er] -> O-phosphoethanolamine[er] + YDR437W_GPI_G6[er]
catalyst: sec_Ted1p_complex

reaction id: YDR437W_GPIRIB_sec_GPIR_complex
reaction equation: 6-O-2-O-alpha-D-mannosyl-(1-2)-{alpha-D-mannosyl-2-O-((2-aminoethyl)phosphoryl)-(1-2)-alpha-D-mannosyl-(1-6)-2-O-((2-aminoethyl)phosphoryl)-alpha-D-mannosyl-(1-4)-alpha-D-glucosaminyl}-O-inositol-P-ceramide C (C26)[er] + YDR437W[er] -> H2O[er] + YDR437W_GPI_G6[er]
catalyst: sec_GPIR_complex

3.3.4 ER O-glycosylation

In yeast, O-glycosylation is initiated in the ER and extended in Golgi. In this section, we describe how the ER O-glycosylation was formulated in the model. There are more than six proteins that are responsible for protein manosylation in yeast. Those proteins make up the complex PMTC, protein O-mannosyltransferase, which transfers mannose residues from dolichyl phosphate D-mannose to protein Ser/Thre residues[21].

In the model, this process was formulated into one reaction:

1. ER O-glycosylation

Example of ER O-glycosylation of YJL137C

| |
|---|
| reaction id:<br>YJL137C_OG_EROG_sec_Pmt2p_Pmt5p_Pmt1p_Pmt6p_Pmt4p_Pmt3p_complex<br>reaction equation: 3 dolichyl D-mannosyl phosphate[er] + YJL137C[er] -> 3 dolichyl phosphate[er] + YJL137C_OG_M1[er]<br>catalyst: sec_Pmt2p_Pmt5p_Pmt1p_Pmt6p_Pmt4p_Pmt3p_complex |

3.3.5 ER N-glycosylation

N-glycosylation is one of the most abundant post-translational modifications in yeast. The first step of N-glycosylation is to transfer the oligosaccharide precursor $Glc_3Man_9GlcNAc_2$ to the protein, which is mediated by the OSTC complex[22]. After that, three glucose residues in the glycan are trimmed by Cwh41[23] and Rot2[22]. Then one of the mannose residues added by Alg9 from N-linked core oligosaccharides is further trimmed by Mns1[22]. This is the last trimming reaction that occurs in the ER before mature proteins migrate to the Golgi. The oligosaccharide precursor synthesis was added to the model in the previous Yeast8 modification step.

This process was divided into five steps in the model.

1. OSTC_complex ER N-glycan transfer
2. ER Glycan trimming I
3. ER Glycan trimming II

4. ER Glycan trimming III
5. ER demanosylation I

Example: ER N-glycosylation for YJL139C

| |
|---|
| reaction id: YJL139C_ERNG_NG_sec_OSTC_complex<br>reaction equation: 5 Glucose(3)Mannose(9)GlucoseNAc(2)-PP-dolichol[er] + YJL139C[er] -> 5 dolichyl phosphate[er] + YJL139C_G3M9[er]<br>catalyst: sec_OSTC_complex<br><br>reaction id: YJL139C_ERNG_FLI_NG_sec_Cwh41p_complex<br>reaction equation: 5 H2O[er] + YJL139C_G3M9[er] -> 5 D-glucose[er] + YJL139C_G2M9[er]<br>catalyst: sec_Cwh41p_complex<br><br>reaction id: YJL139C_ERNG_FLII_NG_sec_Rot2p_complex<br>reaction equation: 5 H2O[er] + YJL139C_G2M9[er] -> 5 D-glucose[er] + YJL139C_G1M9[er]<br>catalyst: sec_Rot2p_complex<br><br>reaction id: YJL139C_ERNG_FLIII_NG_sec_Rot2p_complex<br>reaction equation: 5 H2O[er] + YJL139C_G1M9[er] -> 5 D-glucose[er] + YJL139C_M9[er]<br>catalyst: sec_Rot2p_complex<br><br>reaction id: YJL139C_ERNG_FLIV_NG_sec_Mns1p_complex<br>reaction equation: 5 H2O[er] + YJL139C_M9[er] -> 5 D-mannose[er] + YJL139C_M8[er]<br>catalyst: sec_Mns1p_complex |

##### 3.3.6 Misfolding and ERAD

Protein misfolding is a common cellular process that can produce intrinsically harmful products. In order to reduce the risk, the cell developed a highly efficient system for protein quality control and endoplasmic reticulum-associated degradation (ERAD) for the degradation of misfolded proteins. The ubiquitin-proteasome system (UPS) is a pathway in the cell responsible for the degradation of proteins. The pathway can split into two main processes: ubiquitination and degradation. The ubiquitination process requires three enzymes: E1, E2 and E3[24]. The activating enzyme E1 activates a ubiquitin with one ATP molecule. The activated ubiquitin conjugates enzyme E2. The ubiquitin transfers to the ligase E3 from E2 and then to the misfolded protein when E3 binds to the misfolded protein targeted for degradation. The process repeats itself until the misfolded protein acquires a chain of ubiquitin at least four ubiquitin long. Then, the tagged misfolded protein can then be released from E3 and recognized by the proteasome. In yeast, there are mainly three ERAD pathways: ERAD-C, ERAD-L and ERAD-M, they differ in the E3 ligase part. Both ERAD-L and ERAD-M uses Hrd1 ubiquitin ligase complex (Hrd1, Hrd3p and Der1), but luminal factor Yos9 seems dispensable for ERAD-M. ERAD-C uses the Doa10 ubiquitin ligase complex[25,26]. All protein modifications such as disulfide bond, N-glycosylation and O-glycosylation are reversed in ERAD pathways. We added specific reactions for those processes as the first several reactions in the ERAD reaction, respectively. In order to represent the accumulation of misfolded protein, we added two extra reactions to reflect the occupation of misfolded protein with Kar2 and Pdi1[27].

The misfolding and ERAD is divided into reactions:

1. add Kar2 to the misfolded proteins
2. break the disulfide bond
3. trim one mannose off from the glycan
4. trim the GPI anchor
5. ERAD E3 ligase
6. ERAD ubiquitination
7. trim the glycan
8. misfolding degradation
9. misfolding accumulation with occupation of Pdi1
10. misfolding accumulation with occupation of Kar2

Example of misfolding and ERAD. Since different proteins adopt different reactions based on their protein properties, therefore several proteins are used in this example.

| reaction id: YJL139C_ERAD_sec_Kar2p_complex |
| --- |

reaction equation: 11 H2O[er] + 11 ATP[er] + YJL139C_M9[er] -> 11 H+[er] + 11 phosphate[er] + 11 ADP[er] + YJL139C_M9_misf[er]
catalyst: sec_Kar2p_complex

reaction id: YJR104C_ERAD2A_sec_Pdi1p_complex
reaction equation: 4 glutathione[er] + YJR104C_DSB_misf[er] -> 4 H+[er] + 2 glutathione disulfide[er] + YJR104C_DSB_misf_G1[er]
catalyst: sec_Pdi1p_complex

reaction id: YJL139C_ERAD2B
reaction equation: YJL139C_M9_misf[er] -> YJL139C_M9_misf_G1[er]
catalyst: -

reaction id: YJL139C_ERAD3A_sec_Mns1p_complex
reaction equation: 5 H2O[er] + YJL139C_M9_misf_G1[er] -> 5 D-mannose[er] + YJL139C_M9_misf_G2[er]
catalyst: sec_Mns1p_complex

reaction id: YJR104C_ERAD3B
reaction equation: YJR104C_DSB_misf_G1[er] -> YJR104C_DSB_misf_G2[er]
catalyst: -

reaction id: YJL139C_ERAD4A_sec_Mnl1p_Pdi1p_complex
reaction equation: 5 H2O[er] + YJL139C_M9_misf_G2[er] -> 5 D-mannose[er] + YJL139C_M9_misf_G3[er]
catalyst: sec_Mnl1p_Pdi1p_complex

reaction id: YJR104C_ERAD4B
reaction equation: YJR104C_DSB_misf_G2[er] -> YJR104C_DSB_misf_G3[er]
catalyst: -

reaction id: YKL165C_ERAD5A
reaction equation: H2O[er] + YKL165C_GPI_G6_M9_misf_G3[er] -> 6-O-2-O-alpha-D-mannosyl-(1-2)-{alpha-D-mannosyl-2-O-((2-aminoethyl)phosphoryl)-(1-2)-alpha-D-mannosyl-(1-6)-2-O-((2-aminoethyl)phosphoryl)-alpha-D-mannosyl-(1-4)-alpha-D-glucosaminyl}-O-inositol-P-ceramide C (C26)[er] + YKL165C_GPI_G6_M9_misf_G4[er]
catalyst: -

reaction id: YJL139C_ERAD5B
reaction equation: YJL139C_M9_misf_G3[er] -> YJL139C_M9_misf_G4[er]
catalyst: -

reaction id: YJL139C_ERADL_sec_Cue1p_Ubc6p_Ubc7p_Yos9p_Hrd1p_Hrd3p_Der1p_Usa1p _complex
reaction equation: YJL139C_M9_misf_G4[er] -> YJL139C_M9_misf_G5[er]
catalyst: sec_Cue1p_Ubc6p_Ubc7p_Yos9p_Hrd1p_Hrd3p_Der1p_Usa1p_complex

reaction id: YJL139C_ERADL_sec_Sbh1p_Sss1p_Ssh1p_Cdc48p_Ubx2p_Ufd1p_Npl4p_complex
reaction equation: 8 Ubiquitin_for_Transfer[c] + YJL139C_M9_misf_G5[er] -> 8 Ubiquitin[c] + YJL139C_M9_misf_G6[c]
catalyst: sec_Sbh1p_Sss1p_Ssh1p_Cdc48p_Ubx2p_Ufd1p_Npl4p_complex

reaction id: YKL165C_ERADM_sec_Cue1p_Ubc6p_Ubc7p_Hrd1p_Hrd3p_Der1p_complex
reaction equation: YKL165C_GPI_G6_M9_misf_G4[er] -> YKL165C_GPI_G6_M9_misf_G5[er]
catalyst: sec_Cue1p_Ubc6p_Ubc7p_Hrd1p_Hrd3p_Der1p_complex

reaction id: YKL165C_ERADM_sec_Sbh1p_Sss1p_Ssh1p_Cdc48p_Ubx2p_Ufd1p_Npl4p_complex
reaction equation: 8 Ubiquitin_for_Transfer[c] + YKL165C_GPI_G6_M9_misf_G5[er] -> 8 Ubiquitin[c] + YKL165C_GPI_G6_M9_misf_G6[c]

catalyst: sec_Sbh1p_Sss1p_Ssh1p_Cdc48p_Ubx2p_Ufd1p_Npl4p_complex

reaction id: YNL038W_ERADC_sec_Cue1p_Ubc6p_Ubc7p_Doa10p_complex
reaction equation:
YNL038W_GPI_G6_misf_G4[er] -> YNL038W_GPI_G6_misf_G5[er]
catalyst: sec_Cue1p_Ubc6p_Ubc7p_Doa10p_complex

reaction id:
YNL038W_ERADC_sec_Sbh1p_Sss1p_Ssh1p_Cdc48p_Ubx2p_Ufd1p_Npl4p_complex
reaction equation:
8 Ubiquitin_for_Transfer[c] + YNL038W_GPI_G6_misf_G5[er] -> 8 Ubiquitin[c] + YNL038W_GPI_G6_misf_G6[c]

catalyst: sec_Sbh1p_Sss1p_Ssh1p_Cdc48p_Ubx2p_Ufd1p_Npl4p_complex

reaction id: YJL139C_ERAD7A_sec_Dsk2p_Rad23p_Png1p_Uba1p_complex

reaction equation: YJL139C_M9_misf_G6[c] -> 10 N-acetyl-alpha-D-glucosamine 1-phosphate[c] + 35 D-mannose[er] + YJL139C_misfolding[c]
catalyst: sec_Dsk2p_Rad23p_Png1p_Uba1p_complex

reaction id: YKR058W_ERAD7B_sec_Dsk2p_Rad23p_Png1p_Uba1p_complex
reaction equation: YKR058W_OG_M1_misf_G6[c] -> 2 D-mannose[er] + YKR058W_misfolding[c]
catalyst: sec_Dsk2p_Rad23p_Png1p_Uba1p_complex

reaction id: YNL038W_ERAD7C_sec_Dsk2p_Rad23p_Uba1p_complex
reaction equation: YNL038W_GPI_G6_misf_G4[er] -> YNL038W_GPI_G6_misf_G5[er]
catalyst: sec_Dsk2p_Rad23p_Uba1p_complex

reaction id: YJL139C_degradation_misfolding_c
reaction equation: YJL139C_misfolding[c] -> YJL139C_subunit[c]
catalyst: -

reaction id: YPL091W_cycle_accumulation_sec_pdi1p_ero1p_complex
reaction equation: 10 oxygen[er] + 20 glutathione[er] + YPL091W_DSB_misf[er] -> 10 glutathione disulfide[er] + 10 hydrogen peroxide[er] + YPL091W_DSB_misf2[er]
catalyst: sec_pdi1p_ero1p_complex

reaction id: YPL091W_cycle_accumulation_sec_acc_Kar2p_complex
reaction equation: 120 H2O[er] + 120 ATP[er] + YPL091W_DSB_misf2[er] -> 120 H+[er] + 120 phosphate[er] + 120 ADP[er] + YPL091W_DSB_misfolding_acc[er]
catalyst: sec_acc_Kar2p_complex

3.3.7 COPII transport

The ER is the first membrane compartment of the secretory pathway and is where secretory and membrane proteins traffic in the "vesicular" mode first occurs. The exit from ER requires a distinct set of coat proteins and accessory factors. The first step is per-budding complex formation. Sar1, Sec23 and Sec24 are common proteins among different cargos and coat formation mechanisms. Selective export of soluble luminal cargo requires specific cargo receptors. Yeast Erv29 is required for efficient packaging of the glycosylated alpha-factor pheromone precursor (gpaf) into COPII vesicles and for efficient secretion of carboxypeptidase Y (CPY)[28]. GPI-anchored proteins require adaptor for cargo selection[29]. Emp24, a yeast p24 family protein is one of these studied proteins, which forms heterotrimeric complex with Erp2, Emp24, and Erv25; have been shown working as adaptors for efficient transport of GPI-anchored cargo[30]. Then the budding complex recruits Sec13-Sec31 heterotetramer, providing the outer layer of the coat[31]. Although Sar1, Sec23, Sec24, Sec13 and Sec31 are necessary and sufficient for vesicle formation, additional factors such as Sec16 and Sed4 are also involved in this process[32]. Through interactions with other COPII proteins, Sec16 is thought to facilitate the assembly of the vesicle coat by stabilizing the pre-budding complex, while Sed4 may regulate the vesicle budding process by inhibiting the GTPase-activating protein (GAP) activity of Sec23.

After budding, vesicles have to move toward the target membrane proceeding through defined and progressive steps: tethering, docking, and fusion. Tethering is mediated by Uso1 and Bug1 or multisubunit complexes. COPII complex would tether to the cis-Golgi membrane through directly binding to Sec23 mediated by the TRAPPI complex, including Bet3, Bet5, Trs31, Trs23, Trs33, Trs20, and Trs85[33]. The process of vesicle docking is mediated by another class of proteins named Rabs, which belong to the superfamily of small Ras-like GTPases. Rabs regulates membrane trafficking through interaction with defined effectors. Ypt1 is a Rab protein required ER to Golgi transport and TRAPPI complex activites Ypt1. Fusion is mediated by SNAREs complex such as Bet1 and Bos1[34].

Here in the model, we divided this process into three steps:

1. COPII pre budding (1A lumen cargo 1B transmembrane cargo 1C GPI anchored cargo)
2. COPII formation
3. COPII tethering docking and fusion

And we formulated three different COPII routes for different types of proteins in the model:

| Protein information | Route |
|---|---|
| Transmembrane proteins | coat_trans_membrane |
| GPI anchored proteins | coat_GPI |
| Other proteins | coat_other |

Since it has alternatives, then we use different proteins to show the example of COPII process. Example of COPII_CPI route for protein YBR004C.

```
reaction id:
YBR004C_COPII_GPI_ERGL1C_sec_Sec12p_Sar1p_Sec23p_Sec24p_Emp24p_Erp1p_Erp2
p_Erv25p_Bos1p_Bet1p_complex
reaction equation: H2O[er] + GTP[er] + YBR004C_GPI_G6_M8[er] -> H+[er] +
phosphate[er] + GDP[er] + YBR004C_GPI_G6_M8_GPI_G1_COP[er] catalyst:
sec_Sec12p_Sar1p_Sec23p_Sec24p_Emp24p_Erp1p_Erp2p_Erv25p_Bos1p_Bet1p_complex
```

reaction id:
YBR004C_COPII_ERGL_sec_Sec13p_Sec31p_Sec16p_Sed4p_Sec5p_Sec17p_complex
reaction equation: YBR004C_GPI_G6_M8_GPI_G1_COP[er] ->
YBR004C_GPI_G6_M8_GPI_G2_COP[c]
catalyst: sec_Sec13p_Sec31p_Sec16p_Sed4p_Sec5p_Sec17p_complex

reaction id:
YBR004C_COPII_ERGL_sec_Ypt1p_Uso1p_bug1p_Bet3p_Bet5p_Trs20p_Trs23p_Trs31p_Trs33p_complex
reaction equation: H2O[c] + GTP[c] + YBR004C_GPI_G6_M8_GPI_G2_COP[c] -> H+[c] + phosphate[c] + GDP[c] + YBR004C_GPI_G6_M8[g]
catalyst: sec_Ypt1p_Uso1p_bug1p_Bet3p_Bet5p_Trs20p_Trs23p_Trs31p_Trs33p_complex

Example of coat_trans_membrane route for protein YBR021W.

reaction id:
YBR021W_COPII_TransM_ERGL1B_sec_Sec12p_Sar1p_Sec23p_Sec24p_Bet1p_Bos1p_complex
reaction equation: H2O[er] + GTP[er] + YBR021W[er] -> H+[er] + phosphate[er] + GDP[er] + YBR021W_COP_coated[er]
catalyst: sec_Sec12p_Sar1p_Sec23p_Sec24p_Bet1p_Bos1p_complex

reaction id:
YBR021W_COPII_ERGL_sec_Sec13p_Sec31p_Sec16p_Sed4p_Sec5p_Sec17p_complex
reaction equation: YBR021W_COP_coated[er] -> YBR021W_COP_coated[c]
catalyst: sec_Sec13p_Sec31p_Sec16p_Sed4p_Sec5p_Sec17p_complex

reaction id:
YBR021W_COPII_ERGL_sec_Ypt1p_Uso1p_bug1p_Bet3p_Bet5p_Trs20p_Trs23p_Trs31p_Trs33p_complex
reaction equation: H2O[c] + GTP[c] + YBR021W_COP_coated[c] -> H+[c] + phosphate[c] + GDP[c] + YBR021W[g]

catalyst: sec_Ypt1p_Uso1p_bug1p_Bet3p_Bet5p_Trs20p_Trs23p_Trs31p_Trs33p_complex

Example for the COPII_other route for protein YBR092C.

reaction id:
YBR092C_COPII_normal_ERGL1A_sec_Sec12p_Sar1p_Sec23p_Sec24p_Erv29p_Bet1p_Bos1p_complex
reaction equation: H2O[er] + GTP[er] + YBR092C_M8[er] -> H+[er] + phosphate[er] + GDP[er] + YBR092C_M8_COP_coated[er]
catalyst: sec_Sec12p_Sar1p_Sec23p_Sec24p_Erv29p_Bet1p_Bos1p_complex

reaction id:
YBR092C_COPII_ERGL_sec_Sec13p_Sec31p_Sec16p_Sed4p_Sec5p_Sec17p_complex
reaction equation: YBR092C_M8_COP_coated[er] -> YBR092C_M8_COP_coated[c]
catalyst: sec_Sec13p_Sec31p_Sec16p_Sed4p_Sec5p_Sec17p_complex

reaction id:
YBR092C_COPII_ERGL_sec_Ypt1p_Uso1p_bug1p_Bet3p_Bet5p_Trs20p_Trs23p_Trs31p_Trs33p_complex
reaction equation: H2O[c] + GTP[c] + YBR092C_M8_COP_coated[c] -> H+[c] + phosphate[c] + GDP[c] + YBR092C_M8[g]
catalyst: sec_Ypt1p_Uso1p_bug1p_Bet3p_Bet5p_Trs20p_Trs23p_Trs31p_Trs33p_complex

3.3.8 Protein and signal peptide degradation

Misfolded proteins are transported into the cytosol by the ERAD pathway. Therefore, we need to add one reaction to degrade those proteins into amino acids catalyzed by the proteosome. Similar reactions would also be added for the cleaved signal peptide. The energy cost for the degradation estimated for eukaryotes are around 0.25-1 ATP/aa (the calculation is based on an average protein length of 467 aa)[35]. We adopted the lowest value of 0.25 ATP/aa as the energetic cost for protein degradation and signal peptide. To be noted here, even though that pre and pro sequence in the leader sequence of recombinant protein degraded in ER and Golgi respectively[36]. Here to simplify this process, full leader sequence was cleaved in ER for degradation.

Example of degradation of YPL091W.

reaction id: r_YPL091W_subunit_degradation

reaction equation: 121 H2O[c] + 121 ATP[c] + YPL091W_subunit[c] -> 121 H+[c] + 121 phosphate[c] + 35 L-glutamate[c] + 10 L-methionine[c] + 33 L-alanine[c] + 11 L-glutamine[c] + 627 ADP[c] + 24 L-aspartate[c] + 43 L-glycine[c] + 18 L-arginine[c] + 28 L-asparagine[c] + 29 L-serine[c] + 5 L-cysteine[c] + 15 L-histidine[c] + 32 L-isoleucine[c] + 14 L-proline[c] + 26 L-threonine[c] + 4 L-tryptophan[c] + 19 L-tyrosine[c] + 34 L-leucine[c] + 42 L-lysine[c] + 17 L-phenylalanine[c] + 45 L-valine[c]

catalyst: Mach_Proteasome_complex

Example of degradation of signal peptide for YAL053W.

reaction id: r_YAL053W_SP_degradation

reaction equation: 7 H2O[c] + 7 ATP[c] + YAL053W_sp[c] -> 7 H+[c] + 7 phosphate[c] + L-methionine[c] + 2 L-alanine[c] + 7 ADP[c] + L-glycine[c] + L-arginine[c] + L-asparagine[c] + L-serine[c] + 3 L-cysteine[c] + L-isoleucine[c] + 3 L-threonine[c] + 4 L-leucine[c] + 3 L-phenylalanine[c] + L-valine[c]

catalyst: Mach_Proteasome_complex

##### 3.3.9 Golgi N-glycosylation

N-glycosylated proteins are further modified in Golgi. Upon their arrival, Och1 adds a mannose moiety to the core N-linked oligosaccharide[37]. After that, the Golgi N-glycosylation modification diverges either to form small core-type oligosaccharides or hyper mannan type[38,39]. Since this kind of data is not available for each protein, we adopted the hyper mannan type in the model, which could be easily modified in the future. As for the hyper mannan type N-glycosylation pathway, a heterodimeric complex M-Pol I, consisting of one copy of Van1 and one copy of Mnn9, is the first enzyme that contributes to polymerization of mannose in the Golgi[40]. M-Pol II contains five subunits Mnn9, Anp1, Mnn10, Mnn11, and Hoc1, which further elongates the polysaccharide mannan chain[40]. The α-1,6-mannose backbone is further modified by the addition of α- 1,2-mannoses by Mnn2 and Mnn5, mannosylphosphate residues by Mnn4p and Mnn6p, and α-1,3-mannoses by Mnn1[41].

We divided this process into four steps in the model:

1. Golgi N-glycosylation I with Och1
2. Golgi N-glycosylation II with MPOI
3. Golgi N-glycosylation II with MPOII
4. Golgi N-glycosylation II with Mnn1p Mnn5p and Mnn2p

Example of Golgi N-glycosylation for protein YJL139C

reaction id: YJL139C_GLNG_Golgi_N_linked_glycosylation_I_sec_Och1p_complex
reaction equation: 5 GDP-alpha-D-mannose[g] + YJL139C_M8[g] -> 5 GDP[g] + YJL139C_M8_GNG_G1[g]
catalyst: sec_Och1p_complex

reaction id: YJL139C_GLNG_Golgi_N_linked_glycosylation_II_sec_MPOLI_complex
reaction equation: 45 GDP-alpha-D-mannose[g] + YJL139C_M8_GNG_G1[g] -> 45 GDP[g] + YJL139C_M8_GNG_G2[g]
catalyst: sec_MPOLI_complex

reaction id: YJL139C_GLNG_Golgi_N_linked_glycosylation_III_sec_MPoLII_complex
reaction equation: 150 GDP-alpha-D-mannose[g] + YJL139C_M8_GNG_G2[g] -> 150 GDP[g] + YJL139C_M8_GNG_G3[g]
catalyst: sec_MPoLII_complex

reaction id: YJL139C_GLNG_Golgi_N_linked_glycosylation_II_sec_Mnn1p_Mnn2p_Mnn5p_complex
reaction equation: 5 GDP-alpha-D-mannose[g] + YJL139C_M8_GNG_G3[g] -> 5 GDP[g] + YJL139C_M8_GNG_G4[g]
catalyst: sec_Mnn1p_Mnn2p_Mnn5p_complex

3.3.10 Golgi O-glycosylation

In contrast to N-glycosylation, O-glycans are synthesized by the stepwise addition of monosaccharides besides the first mannose residue added in ER. As for the mannose addition in Golgi, α1,2-mannosyltransferase, Ktr1, Ktr3 along with the Kre2/Mnt1, participates in the addition of the second mannose residue onto O-linked chains[42]. Kre2 has been known to be the primary enzyme responsible for adding the third mannose onto O-glycans[42]. Ktr1 and Ktr3 are also able to add mannose, although to a lesser extent than Kre2. The α1,3- annosyltransferase Mnn1 attaches the fourth mannose residue in the linear chain of up to five mannose residues.

The process was divided into two steps in the model.

1. O-glycosylation mannose extension kre2_ktr1_ktr3
2. O-glycosylation mannose extension Mnn1

Example for Golgi O-glycosylation of YJL137C

| |
|---|
| reaction id:<br>YJL137C_GLOG_Golgi_O_linked_manosylation_I_sec_Kre2p_ktr1p_ktr3p_complex<br>reaction equation: 9 GDP-alpha-D-mannose[g] + YJL137C_OG_M1[g] -> 9 GDP[g] + YJL137C_OG_M1_GOG_G1[g]<br>catalyst: sec_Kre2p_ktr1p_ktr3p_complex<br><br>reaction id: YJL137C_GLOG_Golgi_O_linked_manosylation_II_sec_Mnn1p_complex<br>reaction equation: 6 GDP-alpha-D-mannose[g] + YJL137C_OG_M1_GOG_G1[g] -> 6 GDP[g] + YJL137C_OG_M1_GOG_G2[g]<br>catalyst: sec_Mnn1p_complex |

3.3.11 Mature

We added a mature reaction in the model to indicate the end of the modification.

Example of mature reaction for YJL137C

| |
|---|
| reaction id: YJL137C_Mature<br>reaction equation: YJL137C_OG_M1_GOG_G2[g] -><br>YJL137C_OG_M1_GOG_G2_mature[g] |

| catalyst: - |
| --- |

3.3.12 Sorting

After Golgi processing, mature proteins are transported to their destinations by different vesicle transport.

As for ER or ER membrane proteins, those proteins are transported back to ER via COPI[43]. The COPI complex comprises an ADP-ribosylation factor, Arf1 and the coatomer (Cop1, Sec26, Sec27, Sec21, Ret2, Sec28, and Ret3). The COPI assembly is initiated by the interaction of Arf1 with Golgi membrane. Arf1 activity is controlled by guanine nucleotide exchange factors (GEF) such as Gea1, Gea2, Sec7 etc. Once the COPI arrives ER, Arf1 is inactivated to release the cargo proteins. The uncoating and fusion are two separate steps in ER-Golgi vesicle transport, these processes were lumped in one reaction for simplicity. By hydrolyzing of the ARF1-GTP, the uncoating starts, and then the uncoated vesicle binds to the Golgi membrane by the t-SNAREs[10].

ALP pathway is one of the known trafficking routes from Golgi to Vacuole. Many proteins involved as detecting (Vps1, Swa2), tethering (ClathrinC, Arf1) and docking (t-SNAREC) of the vesicles ALP pathway have been characterized, including Apm3, Apl6, Aps3 and Apl5[44].

The CPY pathway is the default route to the Vacuole from Golgi. A two-step process using AP complexes. The pathway is named so because it has mainly been studied for the trafficking of carboxypeptidase Y to the vacuole. AP-1 complex vesicles can transfer proteins from the trans-Golgi to the early or late endosome. After this, the AP-3 complex vesicle moves proteins from the Golgi/endosome to Vacuole[45].

As for proteins located in the cell membrane or extracellular matrix, there are two types of exocytotic vesicles from the trans-Golgi called light density secretory vesicles (LDSV) and heavy density secretory vesicles (HDSV) (upon density-based separation experiments). LDSV is known to carry constitutively expressed cell membrane proteins such as Bgl2, Pma1 and Gas1; and is believed to emerge from the trans-Golgi and transit directly to the cell membrane. HDSV package soluble, secreted proteins, such as invertase (Suc2) and acid phosphates (Pho11, Pho12, Pho5) which are transcriptionally regulated and induced under certain conditions[46,47].

| Destination | Pathway |
|---|---|
| ERM | COPI |
| ER | COPI |
| VM | ALP |
| V | CPY |
| Cell membrane | LDSV |
| extracellular | HDSV |
| Other | General |

Example of the COPI for protein YJL196C

reaction id:
YJL196C_GLER_COPI_formation_sec_Arf1p_Gea1p_Gea2p_Rer1p_Erd2p_Cop1p_Sec26p_Sec27p_Sec21p_Ret2p_Sec28p_Ret3p_complex
reaction equation: 2 H2O[c] + 2 GTP[c] + YJL196C_mature[g] -> 2 H+[c] + 2 phosphate[c] + 2 GDP[c] + YJL196C_mature_COPI_G1[c]
catalyst:
sec_Arf1p_Gea1p_Gea2p_Rer1p_Erd2p_Cop1p_Sec26p_Sec27p_Sec21p_Ret2p_Sec28p_Ret3p_complex

reaction id:
YJL196C_GLER_COPI_uncoating_and_fission_sec_Rer1p_Ret2p_Cop1p_Sec27p_Sec21p_Bet1p_complex
reaction equation: YJL196C_mature_COPI_G1[c] -> YJL196C_mature[er]
catalyst: sec_Rer1p_Ret2p_Cop1p_Sec27p_Sec21p_Bet1p_complex

Example of the ALP pathway for vacuole membrane protein YJR001W

reaction id:
YJR001W_ALPtransport_sec_Apl6p_Aps3p_Apm3p_Apl5p_Vam3p_Clc1p_Chc1p_Arf1p_Swa2p_Vps1p_complex
reaction equation: 4 H2O[c] + 4 GTP[c] + YJR001W_mature[g] -> 4 H+[c] + 4 phosphate[c] + 4 GDP[c] + YJR001W_folding[vm]
catalyst:
sec_Apl6p_Aps3p_Apm3p_Apl5p_Vam3p_Clc1p_Chc1p_Arf1p_Swa2p_Vps1p_complex

Example of the CPY pathway for protein YKL103C

reaction id:
YKL103C_CPYI_sec_Gga1p_Gga2p_Arf1p_Apl4p_Apl2p_Apm1p_Aps1p_Chc1p_Clc1p_Pep12p_Vps45p_Vps5p_Swa2p_complex
reaction equation: 4 H2O[c] + 4 GTP[c] + YKL103C_M8_GNG_G4_mature[g] -> 4 H+[c] + 4 phosphate[c] + 4 GDP[c] + YKL103C_M8_GNG_G4_mature_CPY_G1[v]
catalyst:
sec_Gga1p_Gga2p_Arf1p_Apl4p_Apl2p_Apm1p_Aps1p_Chc1p_Clc1p_Pep12p_Vps45p_Vps5p_Swa2p_complex

reaction id:
YKL103C_CPYII_sec_Vps4p_Vps27p_Apl6p_Aps3p_Apm3p_Apl5p_Vam3p_complex
reaction equation: H2O[c] + ATP[c] + YKL103C_M8_GNG_G4_mature_CPY_G1[v] -> H+[c] + phosphate[c] + ADP[c] + YKL103C_folding[v]
catalyst: sec_Vps4p_Vps27p_Apl6p_Aps3p_Apm3p_Apl5p_Vam3p_complex

Example of the LDSV for protein YKL217W

reaction id:
YKL217W_LDSV_sec_Arf1p_Sec3p_Sec5p_Sec6p_Sec8p_Sec10p_Sec15p_Exo70p_Exo84p_Sec4p_Chc1p_Clc1p_complex
reaction equation: H2O[c] + GTP[c] + YKL217W_mature[g] -> H+[c] + phosphate[c] + GDP[c] + YKL217W_folding[ce]

catalyst:
sec_Arf1p_Sec3p_Sec5p_Sec6p_Sec8p_Sec10p_Sec15p_Exo70p_Exo84p_Sec4p_Chc1p_Clc1p_complex

Example of HDSV for protein YLR155C

reaction id:
YLR155C_HDSVI_sec_Arf1p_Pep12p_Swa2p_Chc1p_Clc1p_Apl4p_Apl2p_Apm1p_Aps1p_complex
reaction equation: H2O[c] + GTP[c] + YLR155C_M8_GNG_G4_mature[g] -> H+[c] + phosphate[c] + GDP[c] + YLR155C_M8_GNG_G4_mature[ce]
catalyst: sec_Arf1p_Pep12p_Swa2p_Chc1p_Clc1p_Apl4p_Apl2p_Apm1p_Aps1p_complex

reaction id: YLR155C_HDSVII_sec_Vps1p_Chc1p_Clc1p_complex
reaction equation: H2O[c] + GTP[c] + YLR155C_M8_GNG_G4_mature[ce] -> H+[c] + phosphate[c] + GDP[c] + YLR155C_folding[e]
catalyst: sec_Vps1p_Chc1p_Clc1p_complex

Example of general pathway for protein YLR240W

reaction id: YLR240W_transportFromGolgiToOthercompartment
reaction equation: YLR240W_mature_transportFromGolgiToOthercompartment
YLR240W_mature[g] -> YLR240W_folding[g]
catalyst: -

#### 3.4 Enzyme complex formation

In the model, we formulated reactions for enzyme complexes, which serves as catalysts for reactions in metabolism and protein modification processes. If this enzyme contains multiple subunits, then the stichometry for the subunits were included in the equation. Stoichiometries for the subunits are from the PDB database as stated in the previous data collection part as the method in the literature[48].

Example for enzyme complex formation for metabolic reaction r_2141, the complex contains six copies per subunit, thus the stichometry for each subunit is six.

| |
|---|
| reaction id: r_2141_complex_formation |
| reaction equation: 6 YKL182W_folding[c] + 6 YPL231W_folding[c] -> r_2141_complex[c] |
| catalyst: - |

#### 3.5 Complex dilution

The complex dilution reactions in the model are to represent cell division process. Diluted complexes can represent partial protein content in the biomass. Dilution reactions were added for all complexes rather for separate subunits.

Example for complex dilution of r_2141_complex

| |
|---|
| reaction id: r_2141_complex_dilution |
| reaction equation: r_2141_complex[c] -> |
| catalyst: - |

### 4 Turnover rates in the model

#### 4.1 Turnover rates for metabolic complexes

The $k_{cat}$ values for metabolic reactions were acquired from the BRENDA database by matching the EC numbers. The $k_{cat}$ extraction process used the criteria as follows[48]. The $k_{cat}$ values for all organisms were downloaded from BRENDA, with only wildtype enzymes, and only the maximal values for the multiple measurements are kept. The assignment of $k_{cat}$ relies on the matching EC number of the reaction to the dataset. The following criteria was used to determine the $k_{cat}$:

a) $k_{cat}$ values for both substrate and organism matched in the dataset were prioritized.

b) if there is not such fully matched data, then the median of all $k_{cat}$ values within the organism from the same EC number was used.

c) if only substrate was matched but not organism, then the median of all $k_{cat}$ values with matched substrate from the same EC number was used.

d) if neither organism nor substrate was matched, then the median of all available values within the EC number was assigned.

e) if no $k_{cat}$ value was available for the EC number, then the $k_{cat}$ value was set with the median of all assigned values.

Note that the $k_{cat}$ value should be adjusted based on the protein stoichiometry information, e.g., the $k_{cat}$ value should multiply 2 for a dimer enzyme, and the minimal $k_{cat}$ value among subunits was selected for a complex when its subunits had various $k_{cat}$ values. We also manually collect several $k_{cat}$ values, which are available in the GitHub repository: https://github.com/SysBioChalmers/pcSecYeast/tree/main/ComplementaryData/manual_update.xlsx. Besides all these steps, for enzymes with available *in vivo* $k_{cat}$ values ($k_{max}$), we updated the $k_{cat}$ values in the model to *in vivo* $k_{max}$ since it was demonstrated that utilization of *in vivo* $k_{max}$ could improve the model prediction[49]. Functions: `collectkcats`, `updatekcats` and `matchkappToKcat` were used to collect this information and perform changes. Check the corresponding lines in the main function `buildModel` for detailed information.

#### 4.2 Turnover rates for secretory complexes

To get $k_{cat}$ parameters for secretory machinery enzymes, proteome abundance data in PaxDb database[50] combining PSIM mentioned in previous section were used to deduct apparent kinetic parameters. The calculation method is based on the machinery abundance and total abundance of proteins processed by the machinery (Fig. 1). Based on the equation, we can calculate the apparent $k_{cat}$ values. We used the number calculated as the $k_{cat}$ values for secretory machinery complexes, respectively. Function: `SimulateSecParam` was used to get the $k_{cat}$ parameters for the secretory machinery complexes.

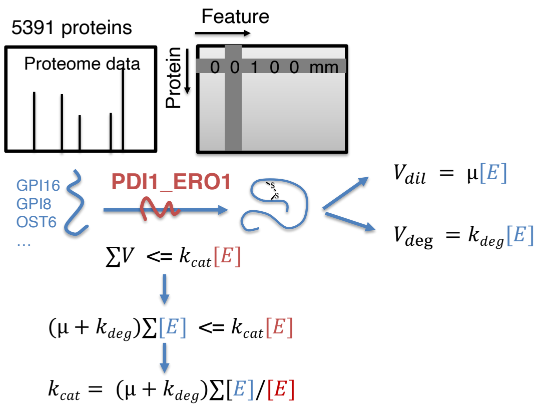

Fig.1 Calculation of secretory machinery $k_{cat}$ parameters.

##### 4.3 Turnover rates for machinery complexes

Ribosome synthesis rate was collected from the literature, which is 2000/min[51]. Considering the ATP turnover for the proteasome is 110/min (bionumber: 109936)[52], while the ATP cost for eukaryote protein degradation is around 100-200 ATP( bionumber: 112155)[35], and we therefore assumed the proteasome catalytic rate to be ~1 protein/min. Given that the average protein length is 467aa, we assumed the proteosome catalytical rate to be 467 aa/min.

The calculation of ribosomal catalytic rate follows the same method done for *Escherichia coli*[53] and *L. lactis*[9]. We assumed the equation of ribosomal catalytic rate follow the Michaelis-Menten-type. We collected the mRNA content, protein content and specific growth rates (Fig. 2)[54–56].

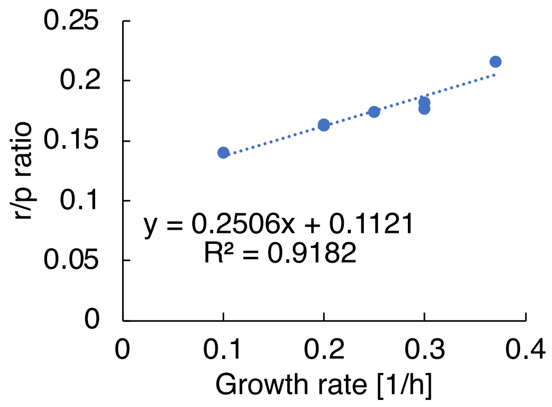

Fig. 2 Ribosome/protein ratio correlates with growth rates.

This study[57] showed that there was a linear correlation between the specific growth rate and RNA/protein ratio in *S. cerevisiae.* From the Fig. 2, we also found clear correlation with the slope being 0.2506 while the intercept 0.1121. The RNA-to-Protein ratio follows the equation.

$$\frac{R}{P} = \frac{\mu}{k_t} + r_0 \quad (1)$$

in which $R$ is total cellular RNA mass (g/gCDW), $P$ is total cellular protein mass (g/gCDW).

Accordingly, we can estimate $k_t$ = 1/0.252 = 3.9904, while $r_0$= 0.1121 based on the equation. Ribosomal catalytic rate (aa/ribosome/s) can be formulated as

$$k_{ribo} = \frac{P_s}{n_r} = \frac{\mu P/m_{aa}}{Rf_{rRNA}/m_{rr}} \quad (2)$$

in which $P_s$ is protein synthesis rate (aa/s), $n_r$ is number of ribosomes, $m_{aa}$is the molecular weight of average amino acid (g/mol), $m_{rr}$ is the mass of rRNA per ribosome (g/mol ribosome), $f_{rRNA}$ is the fraction of rRNA in total RNA.

Using the R/P equation, then

$$k_{ribo} = \frac{\mu/m_{aa}}{f_{rRNA}/m_{rr}} * \frac{k_t * \mu}{k_t * r_0 + \mu} = \frac{V_{max} * \mu}{\mu + K_m} \quad (3)$$

in which:

$$V_{max} = \frac{\mu/m_{aa}}{f_{rRNA}/m_{rr}} * k_t \quad (4)$$

$$K_m = k_t * r_0$$

Given that $m_{aa}$ = 109g/mol, $m_{rr}$ = 1.90E6, $f_{rRNA}$= 0.8 (bionumber: 105192, 100258)[58,59], we can calculate that $V_{max}$ = 24.2, and $K_m$ = 0.447. The ribosomal catalytic rate(aa/ribosome/s) is hence:

$$k_{ribo} = \frac{24.2 * \mu}{\mu + 0.447} \quad (5)$$

#### 5. Constraints

Constraints are required to perform simulations. All constraints were formatted into a linear programming (LP) file for solving as required by the solver SoPlex. Besides the basic flux balanced analysis (FBA) simulation constraint such as:

$$S * V = 0 \quad (6)$$

$$lb_i \leq V_i \leq ub_i \quad (7)$$

which exists in the basic GEM to represent the steady state and the flux for each reaction should be between the lower bound and upper bound.

Constraints to couple metabolic reactions and the corresponding enzymes were also included, which is represented as the rate of a metabolic reaction is constrained by the concentration of the enzyme that catalyzes it:

$$V_{met,i} \leq k_{cat,i} \cdot [E]_i \quad (8)$$

where

$$V_{syn} = V_{dil} \; (9)$$

$$V_{dil} = \mu * [E] \; (10)$$

$$[E] = \frac{V_{syn}}{\mu} \quad (11)$$

In which, $V_{syn}$ represents the formation rate for one enzyme complex, while $V_{dil}$ represents the dilution rate. Dilution rate of one enzyme complex is coupled to the growth rate, thus we can calculate the enzyme abundance from the synthesis rate and the growth rate (eq. 11). Combining together (8-11), we can couple the reaction flux for metabolic reaction with its corresponding enzyme complex formation as:

$$V_{met,i} \leq \frac{k_{cat,i}}{\mu} \cdot V_{syn} \; (12)$$

This type of inequality constraint has been applied to other reactions in protein biosynthesis process, including coupling translation rate and ribosome synthesis rate:

$$\sum V_{trans,i} \leq \frac{k_{cat,translate}}{\mu} \cdot V_{syn,ribosome} \quad (13)$$

coupling ribosome synthesis rate and ribosome assembly rate:

$$V_{ribosome} \leq \frac{k_{cat,ribo\_assembly}}{\mu} \cdot V_{syn,ribo\_assembly} \quad (14)$$

coupling protein degradation rate and proteasome synthesis rate:

$$\sum V_{subunit\_deg,i} \leq \frac{k_{cat,proteasome}}{\mu} \cdot V_{proteasome} \quad (15)$$

coupling post-translational modification rate with corresponding enzyme synthesis rate:

$$\sum V_{\sec\_modification} \leq \frac{k_{cat,i}}{\mu} \cdot V_{\mathrm{syn,sec}\,i} \quad (16)$$

Total proteome constraint is constrained as eq. 17, in which 0.46g is the protein content of 1g biomass.

$$\sum \frac{V_{\mathrm{syn,complex}\,i}}{\mu} + \frac{V_{\mathrm{dummy}}}{\mu} + \frac{V_{\mathrm{dummyER}}}{\mu} + \text{protein content in biomass} = 0.46\ \mathrm{g}\ (17)$$

Besides that, we also included extra constraints in the parameter sensitivity analysis part for CPY accumulation simulation (eq.19-21). Since the protein volume can be roughly considered as the linear correlation with the protein mass[60]. Thus, we can transfer the volume into simple abundance constraint. Therefore, we calculated the maximum value (0.0786g/gCDW) for total ER proteins from multiple available proteome data for *S. cerevisiae* under diverse conditions[1,61–65]. Then, we used this value to constrain the ER protein abundance, which is represented as the sum of each ER protein abundance and then converted to the protein synthesis rate according to the eq. 11.

$$\text{ER volume constraint: } \sum \frac{V_{\mathrm{syn,ER,i}}}{\mu} \leq 0.0786\ (19)$$

The similar constraint was added for ER membrane proteins, ERAD pathway proteins, secretory machinery proteins and retro-translocation enzymes.

$$ER\ membrane\ constraint: \sum \frac{V_{\mathrm{syn,ERM,i}}}{\mu} \leq 0.008\ (18)$$

$$ERAD\ constraint: \sum \frac{V_{\mathrm{syn,ERAD,i}}}{\mu} \leq 0.0125\ (19)$$

$$Sectory\ machinery\ constraint: \sum \frac{V_{\mathrm{syn,SEC,i}}}{\mu} \leq 0.0244\ (20)$$

$$retro - tranlocation\ enzymes\ constraint: \sum \frac{V_{\mathrm{syn,retro,i}}}{\mu} \leq 5.08\text{e-}4\ (21)$$
